# High-resolution scRNA-seq reveals genomic determinants of antigen expression hierarchy in African Trypanosomes

**DOI:** 10.1101/2024.03.22.586247

**Authors:** Kirsty R. McWilliam, Zhibek Keneskhanova, Raúl O. Cosentino, Atai Dobrynin, Jaclyn E. Smith, Ines Subota, Monica R. Mugnier, Maria Colomé-Tatché, T. Nicolai Siegel

**Affiliations:** Division of Experimental Parasitology, Faculty of Veterinary Medicine, Ludwig-Maximilians-Universität München, Munich, Germany; Biomedical Center, Division of Physiological Chemistry, Faculty of Medicine, Ludwig-Maximilians-Universität München, Munich, Germany; School of Biological Sciences, Ashworth Laboratories, Institute for Immunology and Infection Research, University of Edinburgh, Edinburgh, United Kingdom; Institute of Computational Biology, Helmholtz Zentrum München, German Research Center for Environmental Health, Neuherberg, Germany; Department of Molecular Microbiology and Immunology, Johns Hopkins Bloomberg School of Public Health, Baltimore, Maryland, United States of America

## Abstract

Antigenic variation is an immune evasion strategy used by many different pathogens. It involves the periodic, non-random switch in the expression of different antigens throughout an infection. How the observed hierarchy in antigen expression is achieved has remained a mystery. A key challenge in uncovering this process has been the inability to track transcriptome changes and potential genomic rearrangements in individual cells during a switch event. Here, we report the establishment of a highly sensitive single-cell RNA-seq (scRNA-seq) approach for the model protozoan parasite *Trypanosoma brucei*. This approach has revealed genomic rearrangements that occur in individual cells during a switch event. Our data show that following a double-strand break (DSB) in the transcribed antigen-coding gene – an important trigger for antigen switching – the type of repair mechanism and the resultant antigen expression depend on the availability of a homologous repair template in the genome. When such a template was available, repair proceeded through segmental gene conversion, creating new, mosaic antigen-coding genes. Conversely, in the absence of a suitable template, a telomere-adjacent antigen-coding gene from a different part of the genome was activated by break-induced replication. Our results reveal the critical role of available repair sequence in the antigen selection mechanism. Additionally, our study demonstrates the power of highly sensitive scRNA-seq methods in detecting genomic rearrangements that drive transcriptional changes at the single-cell level.

## Introduction

A common strategy used by pathogens to evade the host immune response is antigenic variation. Antigenic variation refers to the ability of a pathogen to systematically alter the expression of an antigen on its surface in order to evade identification and subsequent elimination by the host’s immune response^1^. This strategy is evident in a wide range of evolutionarily distant pathogens, particularly so in *Trypanosoma brucei*^2^*. T. brucei* is a single-celled parasite that lives extracellularly in the bloodstream, adipose tissue^3^, and skin^4^ of its mammalian host and is transmitted by tsetse flies. It is the causative agent of the neglected tropical disease Human African Trypanosomiasis and the wasting disease nagana in both domestic and wild animals. Over 10^7^ identical Variant Surface Glycoproteins (VSG) form a dense, homogeneous surface coat enshrouding the trypanosome. *T. brucei* is able to evade antibody-mediated clearance from the mammalian bloodstream by antigenic variation of this highly immunogenic VSG coat to antigenically distinct VSG isoforms.

The trypanosome genome contains ∼2500 distinct VSG genes, the majority of which are archived in silent subtelomeric arrays in the 11 diploid megabase chromosomes^5^. Most of the remaining VSG genes are located within a highly specialized repertoire of ∼100 30-150kb minichromosomes. Minichromosomes contain a 20-80kb central core of tandem 177bp repeats and one or two VSG genes at their subtelomeres^6,7^. Expression of a VSG relies on the positioning of a VSG gene within one of ∼15 telomere-proximal bloodstream expression sites (BES), which are subject to strict mutually exclusive expression, ensuring that only one is kept transcriptionally active at a time. The general framework of BESs is highly conserved^8,9^. Following an RNA polymerase I promoter, the BES contains up to 11 members of a diverse family of polymorphic genes and pseudogenes termed expression site associated genes (ESAGs). Between the ESAGs and the VSG gene lies a stretch of conserved AT-rich DNA repeats (the so-called ‘70-bp repeats’) often reaching >10kb^10^. Approximately 1.5kb downstream of these repeats is the VSG gene, which itself is followed by >10kb of telomeric TTAGGG repeats. A switch in VSG expression can be facilitated by a transcriptional switch from one BES to another, called an *in situ* switch^11^, or by homologous recombination (HR), whereby a previously silent VSG (either from another BES or from elsewhere in the genomic VSG archive) is recombined into the active BES^12^.

VSG expression follows a semi-predictable and hierarchical pattern, with certain subsets of VSG genes preferentially expressed during specific stages of infection^13,14^. This hierarchy in VSG expression could result from differences in growth rates between cells expressing different VSG isoforms or from differences in the frequency of activation of various VSG genes. Both scenarios are supported by experimental evidence and mathematical models^13–19^. However, existing studies of VSG switching have been limited by the inability to determine VSG expression immediately after a switch in individual cells. Typically, VSG expression is assessed at the population level days after the switch event, making it difficult to distinguish differences in growth rates from differences in activation rates. Consequently, it remains unclear if specific VSG isoforms are preferentially activated and, if so, what mechanism governs this preference during a VSG switch.

To address these questions, we set out to determine the transcriptional profiles of individual cells before, during and after a VSG switch. We developed SL-Smart-seq3xpress, a highly specific and highly sensitive trypanosome-adapted version of the state-of-the-art plate-based Smart-seq3xpress protocol^20^. Using SL-Smart-seq3xpress, we were able not only to determine the newly activated VSGs at the single cell level, but also to determine the switching mechanisms, predict sites of DNA recombination, and reveal the formation of mosaic VSGs.

Our results suggest that following a DSB in the active VSG gene, the presence or absence of a suitable repair template, in the form of a similar VSG gene or pseudogene located elsewhere in the genome, determines what repair mechanism is used and how the new VSG is ‘selected’. In the absence of a repair template, we find that a DSB in the active VSG gene results in one-ended HR within the 70-bp repeats or an upstream ESAG. At the population level, this leads to the reproducible activation of a distinct set of VSG genes previously located immediately adjacent to a telomere in a BES, MES, subtelomeric array or minichromosome. In contrast, when a similar VSG gene or pseudogene exists elsewhere in the genome for use as a repair template, the active VSG gene undergoes segmental gene conversion, resulting in mosaic VSGs.

In summary, our results indicate that the location of a DSB, the presence or absence of a repair template, and the location of the new VSG gene all influence the frequency with which a specific VSG gene is activated. Furthermore, our data suggest that *T. brucei* possesses a highly efficient homology search mechanism that identifies suitable genomic regions to repair DSBs in the active VSG.

## Results

### A Cas9-induced DNA DSB can trigger a switch in VSG expression

Experimentally induced DSBs in both coding and non-coding regions of the actively transcribed BES have been shown to induce VSG switching with various efficiencies depending on the location of the DSB^19,21^. Thus, to investigate the ‘VSG selection mechanism’ controlling the hierarchy of VSG isoform expression, we established a Cas9-based system to induce DSBs at various sites across the active BES. Using a cell line containing a tetracycline-inducible SpCas9 and a phage T7 RNA polymerase capable of transcribing sgRNA molecules from a stably transfected plasmid, our approach is similar to that used in a recently published study^19^.

We designed a sgRNA that directs Cas9 to induce a DSB within the 5’end of the actively transcribed VSG (*VSG-2*, nucleotide position 152, Fig. 1a) and tested the inducibility and efficiency of DSB generation. The ability to efficiently generate DSBs in cells transfected with the *VSG-2* sgRNA was confirmed by western blotting and immunofluorescence analysis (IFA) using an antibody specific to γ-H2A, a marker of DSBs (Fig. 1b,c). Next, to assess the efficiency by which this approach would lead to a switch in VSG expression, we induced Cas9 expression and analyzed VSG-2 expression by flow cytometry using a fluorophore-conjugated VSG-2 antibody at 0, 24, 48, 72 and 96 hours post-induction. Our results showed that the uninduced starting population was largely VSG-2 positive, but that VSG-2 expression was lost from over 94% of the population by 96 hours post-Cas9 induction (Fig. 1d). Sanger sequencing of cDNA isolated from five VSG-2 negative sub-clones confirmed that the cells had switched expression to a new VSG (Supplementary Data 1). Taken together, these results confirm that CRISPR/Cas9 can be used to reliably and efficiently generate a DSB within the CDS of the actively expressed VSG and to trigger a switch in VSG expression.

**Figure 1.**
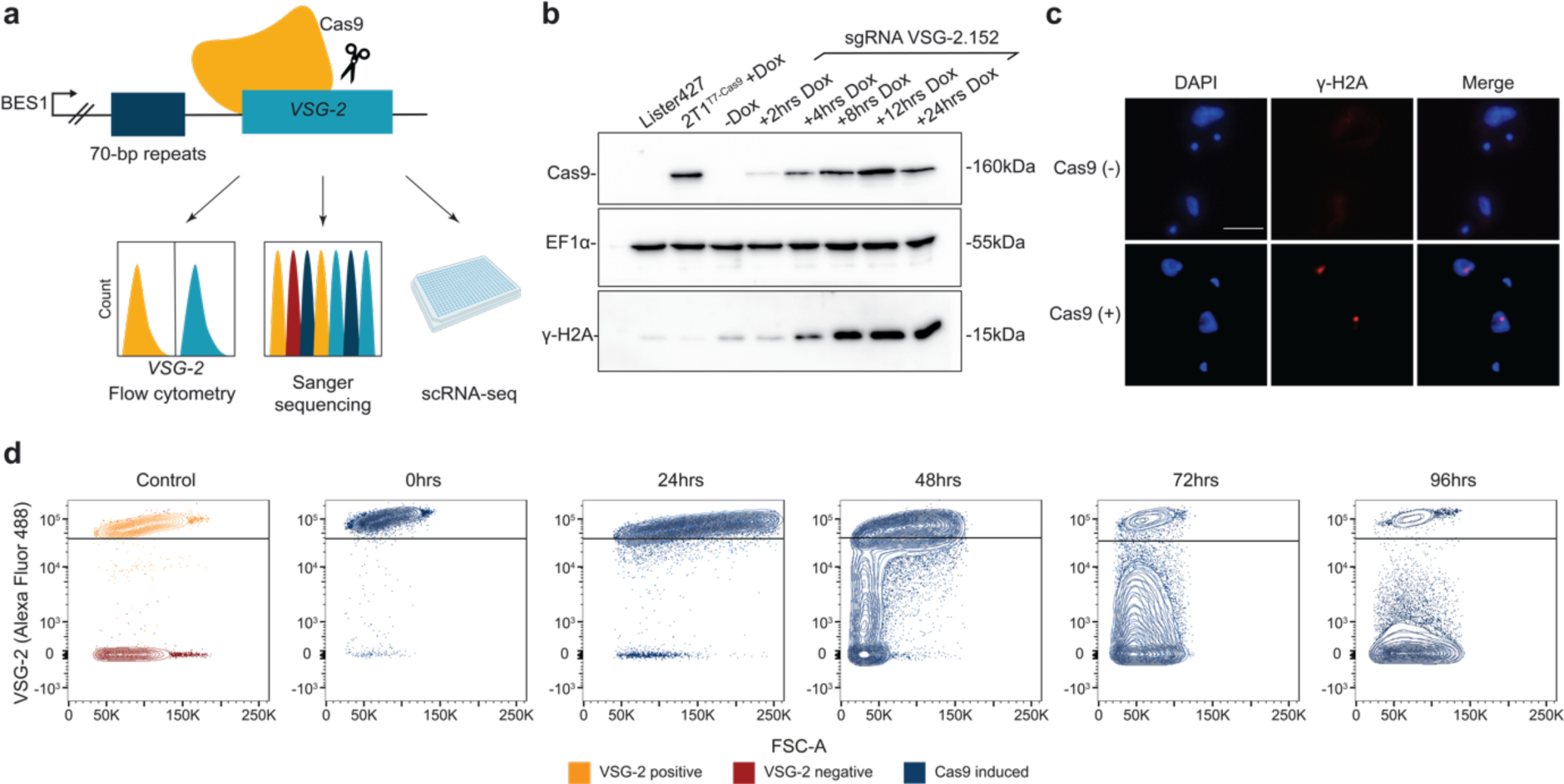
A CRISPR-Cas9 induced DSB in the actively expressed VSG induces a VSG switch. **a,** Strategy to investigate VSG switching and template selection at the single cell level. **b,** Western blot analysis of Cas9 and γ-H2A protein expression in cells capable of doxycycline-inducible expression of Cas9 and a sgRNA targeting nucleotide site 152 of *VSG-2* (sgRNA VSG-2.152). A wild-type cell line (Lister 427) and a cell line not transfected with the VSG-2 sgRNA but capable of inducible Cas9 expression (2T1^T7^ ^Cas9^) served as controls. EF1α served as a loading control. **c,** IFA of γ-H2A expression in cells transfected with sgRNA VSG-2.152 either before (Cas9(-)) or 4 hours after Cas9 induction (Cas9 (+)). γ-H2A was detected using an anti-γ-H2A antibody. Scale bar=10µm. **d,** FACS analysis of VSG-2 expression in sgRNA VSG-2.152 transfected cells in a time course until 96 hours post-Cas9 induction. VSG-2 expression was monitored using a fluorophore-conjugated (Alexa Fluor 488) anti-VSG-2 antibody. At least 10,000 events were analysed per sample. On the left panel, VSG-2 positive and VSG-2 negative cell lines are shown as controls.

### Implementation of SL-Smart-seq3xpress single-cell RNA-sequencing

Next, to determine the transcriptome of individual parasites during a VSG switch, we sought to establish a suitable scRNA-seq approach. In the past, VSG switch studies have typically cloned individual cells after induction of a switch and analyzed the resulting population^22^ or used bulk RNA-seq to analyze VSG expression within a population^19^. However, cloning of individual cells after induction of a VSG switch does not provide information about early time points (some events may result in cell death or a second switch event may occur prior to VSG expression analysis) and is limited by scale. Furthermore, bulk RNA-seq analysis cannot distinguish between monogenic and multigenic VSG expression and does not yield information about VSG switch mechanisms or the site of DNA recombination in individual cells. Therefore, we decided to implement an appropriate scRNA-seq approach. Crucial to our analysis was the availability of a scRNA-seq approach that was a) scalable and allowed multiplexing (allowing us to analyze multiple time points and replicates simultaneously), b) highly sensitive (*T. brucei* has 50 times less mRNA molecules than the average mammalian cell^23,24^ and ESAGs are expressed at low levels), c) highly accurate (allowing us to precisely quantify the degree of mutually exclusive VSG expression without confusing sequencing artefacts for biological truths), d) able to provide full-length transcript information (helping us distinguish between VSG isoforms).

Following our positive experience with the plate-based Smart-seq2 method^10^, we set out to implement a *T. brucei*-adapted version of the most recent Smart-seq protocol: Smart-seq3xpress. Smart-seq3xpress combines full-length transcript detection with a 5’ UMI counting strategy at a nanoliter scale, significantly reducing the cost per cell without sacrificing sensitivity^20,25^. UMIs are used to eliminate PCR amplification bias and are therefore important for accurate transcript level quantification. To take advantage of the fact that all mature mRNA molecules in *T. brucei* contain a conserved 39nt spliced leader (SL) sequence^26^, we omitted the template switch step and instead performed the second-strand cDNA synthesis and amplification using primers annealing to the SL sequence (Fig. 2a). This required us to modify the standard Smart-seq3xpress oligos and therefore the 8bp UMI, 11bp UMI-identifying tag and partial Tn5 motif were moved from the template switch oligo to the oligodT primer (Fig. 2a, Extended Data Fig. 1a). To account for the fact that *T. brucei* cells contain significantly less RNA than the mammalian HEK293FT cells with which the Smart-seq3xpress protocol was developed, we adjusted the transposase and oligodT primer concentrations to maximize transcript diversity and the number of UMI-containing reads without losing the transcript internal reads that do not contain UMI but are important for distinguishing VSG isoforms (Extended Data Fig. 1b,c).

**Figure 2.**
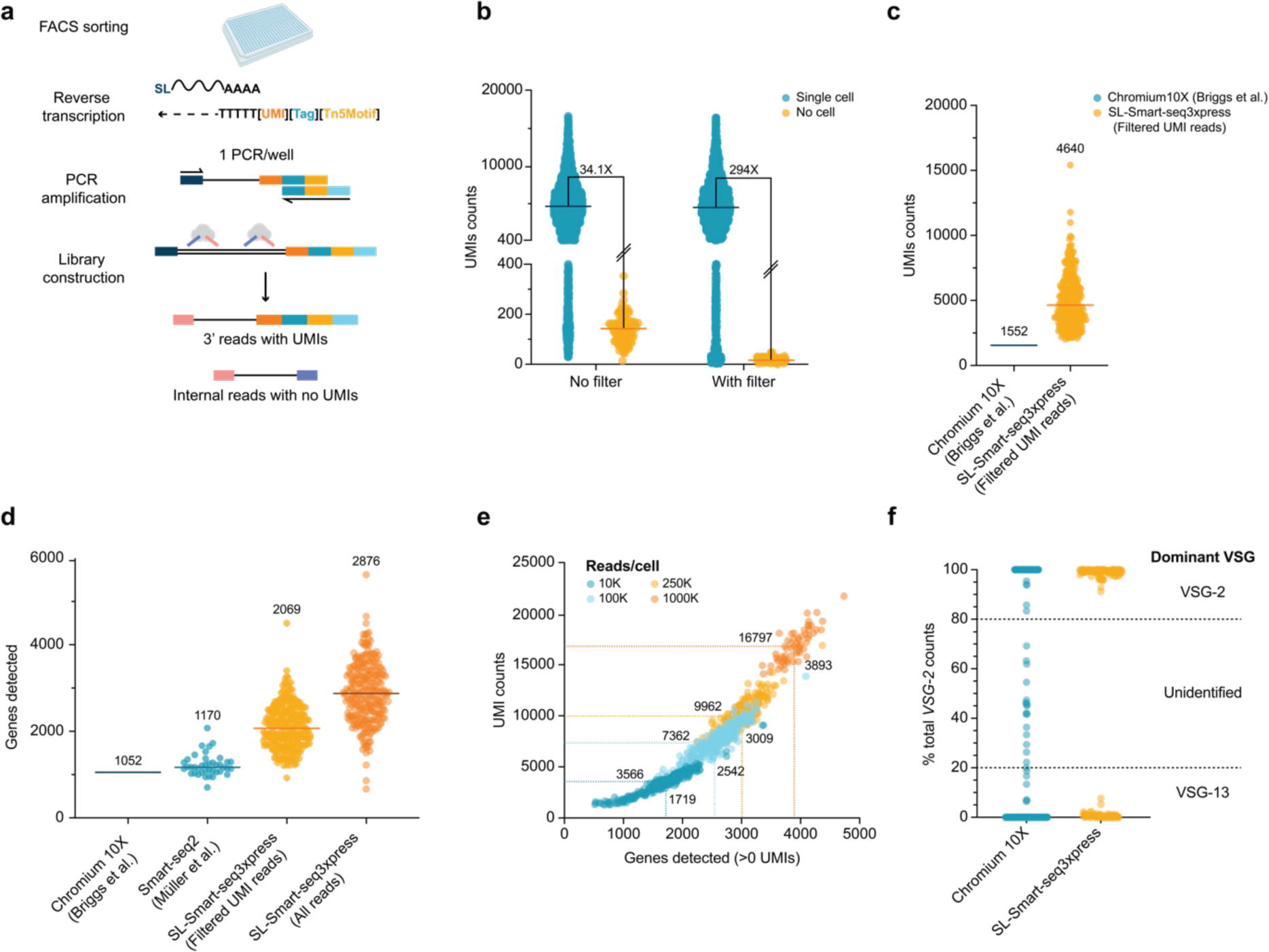
SL-Smart-seq3xpress is a highly sensitive and specific scRNA-seq approach. **a,** Schematic of SL-Smart-seq3xpress library preparation. **b,** Number of UMIs detected by SL-Smart-seq3xpress in single cells (blue) or no cell wells (orange) either without (No filter) or with (With Filter) bioinformatics removal of index hopped reads. The ratio median UMI counts (single cell/no cell) is shown as a number between the two conditions. The median UMI count is shown as a black line. Number of wells analyzed: single cells - 2214, no cells - 120. **c,** Comparison of the median number of UMIs detected by Chromium 10X (data from Briggs et al, 2021^28^) or SL-Smart-seq3xpress. **d,** Comparison of the median number of genes detected by Chromium 10X (data from Briggs et al, 2021^28^), Smart-seq2 (data from Müller et al, 2018^10^) or SL-Smart-seq3xpress. The median UMI/gene count is shown as a number above a dataset and as a bold line. For the Smart-seq2 and SL-Smart-seq3xpress datasets, each dot represents an individual cell. For the Chromium 10X dataset, the line represents the median UMI/gene count from two replicate experiments. Libraries are downsampled to 75k reads/cell. Number of cells analyzed: Chromium 10X - 8599, Smart-seq2 - 40, SL-Smart-seq3xpress - 292. **e,** Detected UMIs vs genes in a SL-Smart-seq3xpress library at increasing read depths. Each single cell is represented by a colored dot. The median gene/UMI count for each read depth is represented by the dotted lines and numbers. **f,** Percentage of *VSG-2* transcript counts, relative to the sum of VSG-2 and VSG-13 transcript counts, in Chromium 10X and SL-Smart-seq3xpress single cell libraries prepared from mixed populations of *VSG-2* expressing (P10 cell line) and *VSG-13* expressing (N50 cell line) cells. The thresholds for defining if a cell is a *VSG-2* or *VSG- 13* expresser (above 80% and below 20%, respectively) are shown by the dotted lines. Total number of cells analyzed: Chromium 10X - 185, SL-Smart-seq3xpress - 185.

Next, we assessed the specificity of our sequencing approach. Following the introduction of patterned flow cells in the latest generation of Illumina sequencers, several groups have reported library index hopping as a significant source of error^27^. Such index hopping can result in the incorrect assignment of reads from one cell to another and would limit our ability to investigate the regulation of mutually exclusively expressed genes, such as the VSG (Extended Data Fig. 1d). To address this challenge, we implemented scSwitchFilter (https://github.com/colomemaria/scSwitchFilter), a), a bioinformatic tool that removes index hopped reads based on the assumption that reads with the correct gene-UMI-index combination should be more abundant than reads with the same gene-UMI combination arising from an index hopping event (Extended Data Fig. 1d). Application of this filtering approach drastically decreased the observed number of index hopping events, with the median UMI count from a single cell now 294-fold higher than that from control wells not containing a cell (Fig. 2b). We therefore applied the index hopping filtering pipeline to all quantitative transcriptome analyses.

### SL-Smart-seq3xpress reliably quantifies VSG transcript levels

Having optimized our SL-Smart-seq3xpress pipeline, we compared the sensitivity (the number of genes and UMIs detected per cell) of SL-Smart-seq3xpress to the best available 3’ Chromium Single Cell (10X Genomics) dataset for *T. brucei*^28^. Since the 10X data was sequenced to ∼75,000 reads per cell, we downsampled our SL-Smart-seq3xpress data to the same number of reads and found it to be significantly more sensitive, detecting a median of 2,876 genes and 4,640 UMIs per cell vs 1,052 genes and 1,552 UMIs detected in the published 10X Genomics dataset (Fig. 2c and Fig. 2d, respectively). At higher sequencing depth (1 million reads per cell sequenced) we were able to detect a median of 16,797 transcripts per cell (Fig. 2e). This corresponds to ∼84% of the predicted total number of mRNA molecules in a single cell^24^. Thus, our SL-Smart-seq3xpress approach ranks among the most sensitive scRNA-seq methods reported^29^.

To test whether SL-Smart-seq3xpress was able to reliably detect VSG transcripts and distinguish between different VSG transcripts originating from different cells, we mixed isogenic cells expressing one of two different VSGs, either *VSG-2* or *VSG-13*^30,31^, in equal proportions and generated sequencing libraries. For comparison, we generated another sequencing library from the same mixture of cells using the 5’ Chromium Pipeline. We defined a cell’s VSG expression as ‘mutually exclusive’ when more than 80% of the cell’s *VSG-2* and *VSG-13* transcripts mapped to only one of the two VSG genes. When we sequenced the mixed population using our SL-Smart-seq3xpress pipeline, we observed clean separation between *VSG-2* and *VSG-13* expressers, with 100% of the cells reaching the defined threshold for mutually exclusive expression, i.e. a relative *VSG-2* fraction above 80% or below 20% (Fig. 2f). For every cell that had a relative *VSG-2* fraction of below 20%, each correspondingly had a *VSG-13* fraction of over 80%. In contrast, when cells from the same mixed population were sequenced using the 5’ Chromium method, 11% of the cells did not surpass the defined threshold (Fig. 2f). Droplet-based single-cell RNA-seq pipelines, such as those used by 10X Genomics, have been reported to suffer from a certain degree of ambient RNA contamination and the presence of cell ‘doublets’ in the drops, and it seems likely that this explains the high proportion of ‘double VSG expressers’ in our 5’ Chromium dataset. Similar observations were made when *T. brucei* and *L. mexicana* cells were mixed during a 3’ Chromium analysis^28^. When we increased the threshold from 80% to 90%, we still found that 100% of the cells sequenced using SL-Smart-seq3xpress show unambiguous mutually exclusive VSG expression (Fig. 2f). For the 5’ Chromium platform, the percentage of cells not meeting the threshold increased from 11% to 14%, similar to that reported in a recent VSG analysis using the 5’ Chromium platform^32^. Our low background is consistent with previous observations suggesting that plate-based scRNA-seq approaches suffer less from ambient RNA contamination than droplet-based approaches^33^. Overall, our SL-Smart-seq3xpress pipeline, combined with the bioinformatic cleanup of index-hopped reads, appears to be very well suited for the analysis of mutually exclusively expressed genes such as VSGs.

### A DSB in the active VSG-2 leads to activation of telomere-adjacent VSGs only

Previous studies of experimentally-induced VSG switches have typically cloned a relatively small number of cells after induction of a DSB and allowed the clones to grow to a sufficient density before analysis. This made it impractical to study the events during or immediately after a DSB-induced switch^19,21,34–36^. However, after establishing our SL-adapted version of Smart-seq3xpress, which was able to detect VSG expression in single cells with high precision and minimal background, we were able to monitor VSG expression in single cells before and at various times after induction of a DSB within *VSG-2*.

To investigate the VSG selection mechanism, we generated a DSB at nucleotide position 152 of the actively expressed *VSG-2* and prepared SL-Smart-seq3xpress sequencing libraries from 369 cells at 0 hours, 24 hours, 48 hours, 72 hours, 96 hours and 10 days after Cas9 induction. This was done for two biological replicates for a total of 738 cells per timepoint. As already observed at the protein level by flow cytometry (Fig. 1d), the scRNA-seq data indicated that the generation of a DSB within *VSG-2* led to a rapid decrease in the proportion of cells that dominantly expressed *VSG-2* (Fig. 3a). In these cells the total number of VSG transcripts dropped by almost 10-fold, pointing to a transcriptional arrest of the active BES (Extended Data Fig. 2), with no other VSG transcripts upregulated. By 48 hours post-Cas9 induction these cells comprised the majority of the population. However, we then began to observe that a broad but well-defined set of VSG genes had been activated at the population level, including VSGs located in BESs, subtelomeric VSG-arrays, minichromosomes, and metacyclic expression sites (MES, expression sites that are active in the infectious metacyclic form prior to BES activation) (Fig. 3a and Fig. 3b). The similarity of the set of activated VSGs between the two replicates was striking, suggesting hard-wired mechanisms leading to the preferential activation of some VSGs over others, as observed previously^19,36^. By day 10 post-induction, most cells expressed a single dominant VSG, and several switched clones, particularly those expressing VSG-9 and VSG-18, had begun to outgrow, suggesting VSG-specific fitness advantages contribute to a reduction in VSG expression heterogeneity within the population (Fig. 3a). Given that these VSGs are amongst the largest in the genome, this observation suggests that VSG growth dynamics are not primarily governed by VSG length, as previously suggested^18^.

**Figure 3.**
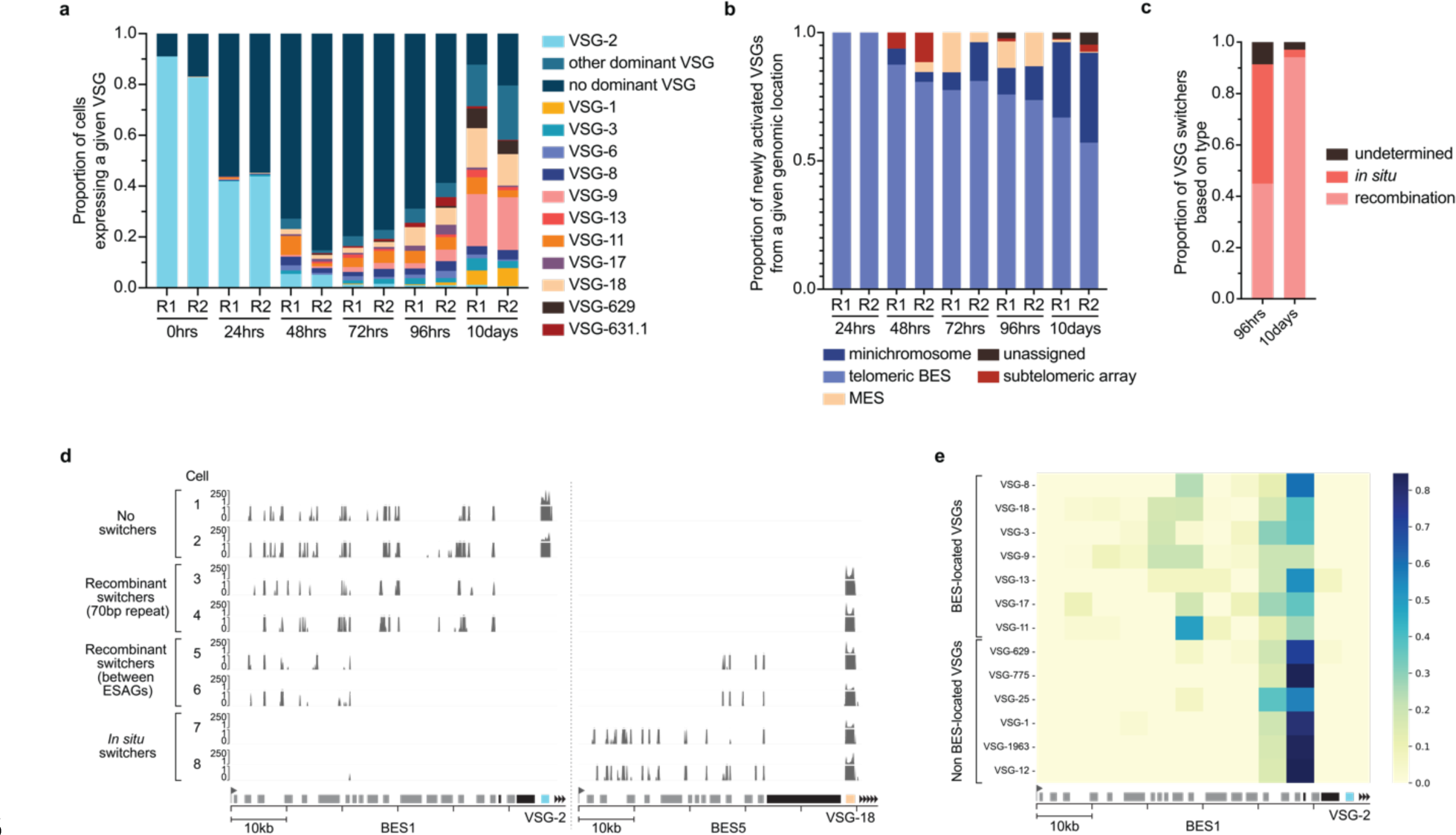
DSBs in the active VSG-2 CDS led to activation of telomere-adjacent VSGs. **a,** VSG expression in single cells before and after the induction of a DSB in the *VSG-2* CDS at nucleotide position 152, as measured by SL-Smart-seq3xpress. Two biological replicate (R) inductions were performed. Total number of cells analyzed per timepoint and replicate are as follows: 0 hours R1, 312; 0 hours R2, 302; 24 hours R1, 308; 24 hours R2, 283; 48 hours R1, 147; 48 hours R2, 271; 72 hours R1, 305; 72 hours R2, 255; 96 hours R1, 289, 96 hours R2, 286; 10 days R1, 336; 10 days R2, 323. **b,** Proportion of cells at each timepoint from **(a)** expressing a new VSG from a given genomic location. Cells that did not clearly express a single VSG (no dominant VSG in **(a)**) or cells that had not switched VSG were not considered for this analysis. If the genomic location of the newly activated VSG was not known, it was termed ‘unassigned’. **c,** Percentage of cells, at 96 hours and 10 days post Cas9 induction, that switched VSG expression by DNA recombination, by *in situ* transcriptional switch, or that could not be determined. **d,** Single cell sequencing read coverage on BES1 and BES5 showcasing different VSG switching scenarios for cells that had switched to *VSG-18* expression following a DSB in the *VSG-2* CDS. The read coverage is shown for 8 cells, 2 cells per VSG switching scenario: no switching (‘No switchers’, cells expressing *VSG-2*), switching by recombination around the 70-bp repeats (‘Recombinant switchers’ (70-bp repeat)), switching by recombination between ESAGs (‘Recombinant switchers’ (between ESAGs)), and *in situ* switching (‘*In situ* switchers’). **e,** Heatmap summarising transcriptional signal end positions on BES1 for recombinant switcher cells after a DSB in the VSG-2 CDS. Each box in the heatmap represents the fraction of cells, for switchers to a given VSG (rows), which transcriptional signal ends at the given 5kb bin of BES1 (columns). VSGs expressed in at least 10 cells were considered.

Significantly, when we considered all newly activated VSGs, a consistent pattern emerged – every newly activated VSG was situated at the terminus of a chromosome, immediately adjacent to a telomere. Out of the activated VSGs, 12 were located in a BES, 5 in a MES, 19 were located on minichromosomes, and 3 were the telomere-proximal VSG gene in a subtelomeric VSG array (Extended Data Table 1). This distinct preference for telomere-adjacent VSGs, despite the presence of hundreds of non-telomeric VSGs in the genome, strongly suggests a preferential use of a specific homologous recombination (HR) mechanism, known as break-induced replication (BIR). This one-end dependent HR mechanism appears to be the primary pathway for VSG expression switching through recombination following a DSB in *VSG-2*.

### Transcription of BES1 confers fitness advantage

To better understand the mechanism governing VSG selection, we next sought to determine whether the initial VSG switch occurred by DNA recombination or by a transcriptional shift to an alternative BES. The high sensitivity of our SL-Smart-seq3xpress approach allowed us to detect transcripts from ESAGs and thus distinguish whether the original BES (BES1) was still transcribed or whether a new BES had been activated (Fig. 3d). At 96 hours post-Cas9 induction, of 169 cells that had switched transcription to an alternative dominant VSG, we found that 46.5% had undergone an *in situ* switch to a different BES, while 44.9% had switched by DNA recombination into BES1 (Fig. 3c). For those cells that had switched by recombination, analysis of ESAG transcripts also allowed us to approximate the sites of DNA recombination at the single cell level. Our data indicated that most recombination events occurred in (or near) the 70-bp repeats upstream of *VSG-2* (Fig. 3e). However, recombination was not restricted to the 70-bp repeats; in particular, for switchers to *VSG-11*, *VSG-9*, *VSG-18* and *VSG-8*, we observe a considerable fraction of cells whose transcriptional profiles suggest recombination between ESAGs (Fig. 3e and Extended Data Fig. 3, left panel).

Interestingly, 10 days after Cas9 induction, we observed a surprising shift in the population composition compared to the 96 hours time point. Now 94.1% of the cells were recombinant switchers transcribing BES1, suggesting that there is a strong fitness advantage in keeping BES1 active (Fig. 3c). Several *in situ* switchers, including those that had switched expression to a MES and were detected 96 hours after Cas9 induction, were completely absent from the population at the later time point.

Several studies have proposed DNA recombination to be the predominant means of VSG switching following induction of a DSB in BES1^21,34,36^. Our data instead shows that *in situ* switches in VSG expression are as likely to occur as DNA recombination-based switches following a DSB in *VSG-2*. However, these variants appeared either to have been outcompeted by cells that had maintained BES1 expression or to have switched back to transcribing BES1 once the DSB was repaired.

### The location of a DSB affects the outcome of its repair

Following our observations that HR was concentrated at specific regions of BES1, and that the repair templates were very consistent between replicates, we next investigated if the position of a DSB affects the position of recombination and with that, the repair template choice. To this end, we generated four cell lines in which Cas9 was used to generate a DSB at either the center of the actively expressed *VSG-2* gene (nucleotide 782); upstream of the 70-bp repeats on BES1 (BES1 nucleotide 40225); between the second set of 70-bp repeats and the *VSG-2* CDS (BES1 nucleotide 54824); or between VSG-2 and the telomere (BES1 nucleotide 58149) (Fig. 4a). Successful DSB generation for all cell lines was confirmed by analysis of γ-H2A signal intensity using western blotting (Extended Data Fig. 4d-h).

**Figure 4.**
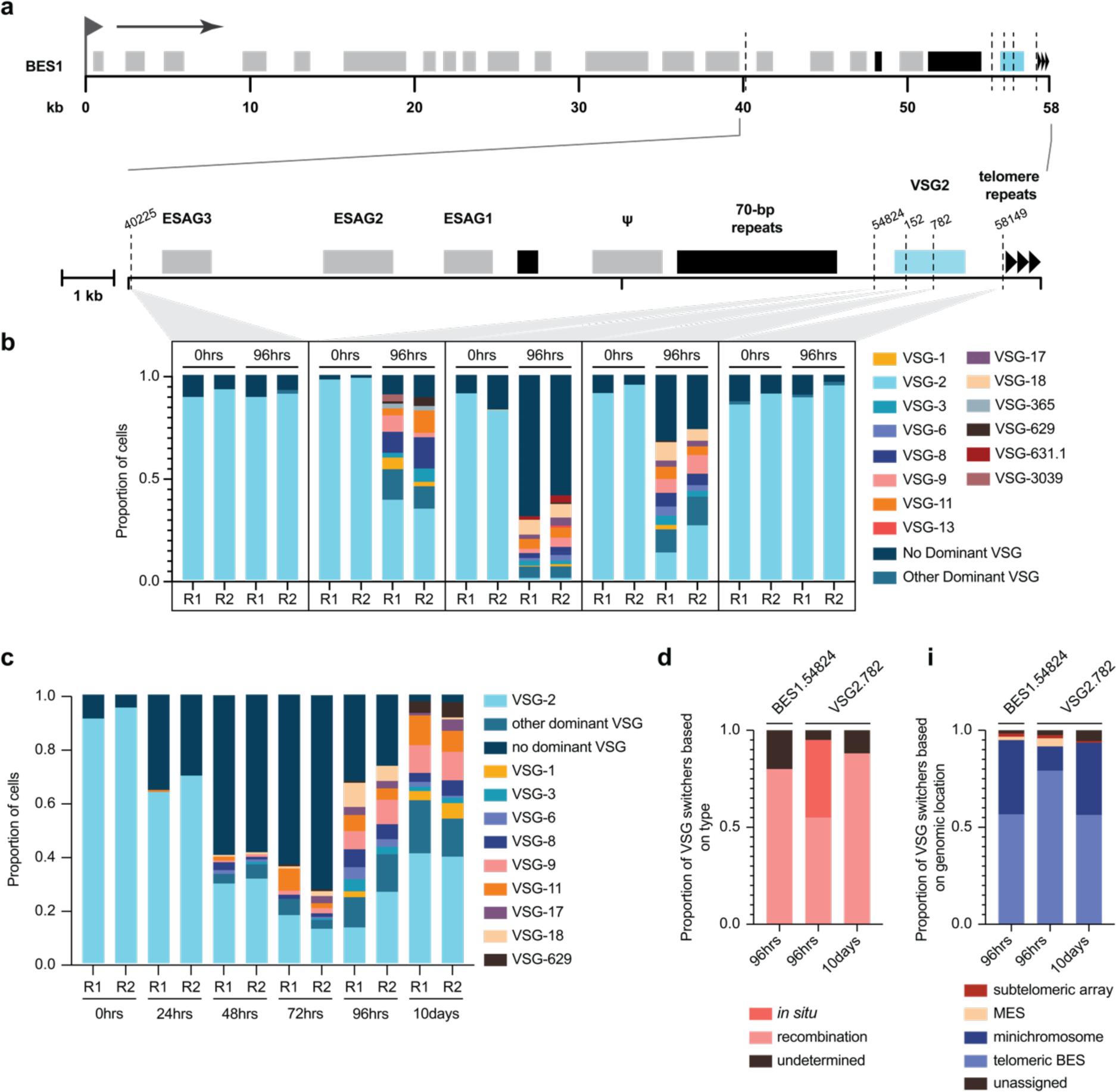
DSB position in the active BES impacts the switching outcome. **a,** A schematic map of BES1 with a zoom-in to the region encompassing the cut sites (dashed lines). The coordinates are relative to the promoter or to the 5’-end of the VSG CDS. **b,** Summary of scRNA-seq results showing the proportion of cells expressing a given VSG for each cut site along the expression site at 96 hours and 10 days post-Cas9 induction. Data is shown for each of two biological replicates (R1 and R2) separately. **c,** scRNA-seq results of the time course experiment following a DSB induction at nucleotide position 782 in the *VSG-2* CDS, showing the proportion of cells expressing a given VSG at each time point for each replicate. **d,** The left plot shows the proportion of cells that had switched VSG expression by recombination or an *in situ* switch of transcription at 96 hours after DSB induction at position 54824 of BES1, and at 96 hours and 10 days time points for the cut site at nucleotide position 782 in the *VSG-2* CDS. Cells for which the switching mechanism could not be clearly determined, are labeled as ‘undetermined’. The right plot shows, for the same datasets, the proportion of switcher cells grouped by the genomic location of the newly activated VSG. Cells expressing a VSG for which the genomic location is not known have been categorized as ‘unassigned’.

Following the induction of a DSB in the middle of *VSG-2* (position 782), we observed a pattern of active VSGs that was remarkably similar to that observed for the cut in the 5’-region of *VSG-2* (position 152) (Fig. 4c). Again, we observed at early time points a significant proportion of cells that did not express a single VSG dominantly, followed by the activation of telomeric VSGs via both *in situ* and DNA recombination-based mechanisms (Fig. 4d).

Next, we targeted Cas9 to sites on BES1 outside of *VSG-2*. When a DSB break was generated just upstream of *VSG-2*, between the 2^nd^ set of 70-bp repeats and *VSG-2*, we observed a moderate induction of VSG switching and again found that the pattern of the preferentially activated VSGs was very similar to that generated by a DSB within *VSG-2* (Fig. 4b). Previous studies of an experimentally induced DSB at this locus have also reported frequent activation of the same VSGs observed in our dataset, such as *VSG-8*, *VSG-11*, *VSG-9*, and *VSG-1*^36^. Visual inspection of the mapped reads confirmed that the alternative VSGs were preferentially activated by DNA recombination around the 70-bp repeats on BES1 (Extended Data Fig. 3, right panel). In stark contrast, when we induced a DSB at the cut-site upstream of the 2^nd^ 70-bp repeats, we observed no switching except in one cell (Fig. 4b). This was despite robust Cas9 induction and elevated γ-H2A signal (Extended Data Fig. 4e,f). We found this particularly interesting because we had previously observed DNA recombination upstream of this cut site when cutting within *VSG-2* (Fig. 3e, Extended Data Fig. 3), demonstrating that upstream DNA recombination is possible. Since this DNA recombination does not happen when we cut upstream of the 70-bp repeats, but only when we cut downstream, we postulate that different mechanisms of DNA repair may be triggered when a DSB occurs either up- or downstream of the 70-bp repeats on the active BES, in line with what has been observed previously^21,36^.

In summary, our analysis of 5 breakpoints along BES1 containing VSG-2 suggests that most recombination events occur at or near the 70-bp repeats. Consistent with this observation, we find that the position of the DSB relative to the 70-bp repeat region is more important in triggering a VSG switch than the sequence immediately flanking the DSB.

### *T*. *brucei* preferentially repairs DSBs in the active VSG by segmental gene conversion

It is important to note that *VSG-2* is unusual among VSG genes, in that its CDS has very little sequence similarity to any other VSG gene in the genome. Therefore, to conclusively address whether local sequence homology influences the identity of the activated VSGs following a DSB, we decided to determine whether the outcome of our switching experiments would change if we cut in an active VSG that had a high degree of sequence similarity to other VSGs in the genome.

To generate cell lines expressing an active VSG gene that shared sequence homology with at least one other VSG gene or pseudogene in the genome, we induced a DSB at position 152 of *VSG-2*, subcloned surviving switchers, determined the actively expressed VSG by Sanger sequencing of the clones’ cDNA, and selected a clone expressing *VSG-8* from BES1 (Extended Data Fig. 5a). *VSG-8* has a second copy in the genome and shares regions of DNA sequence homology with at least 5 other VSG genes and pseudogenes in the genome, as determined by BLAST.

In this *VSG-8* expressing clone, we then induced a DSB in the *VSG-8* CDS (either at nucleotide position 609 or 1105, Fig. 5a) using sgRNAs that were designed to target exclusively *VSG-8*, and not one of the similar VSG genes. Efficient DSB induction was confirmed by analysis of γ-H2A signal intensity using western blotting (Extended Data Fig. 5b). Interestingly, we observed a milder restriction of population growth upon DSB generation at both sites, compared to that observed following the cuts in *VSG-2* (Extended Data Fig. 4a-c). SL-Smart-seq3xpress libraries were prepared from cells before and 96 hours after Cas9 induction for each of the two cut sites for two biological replicates each.

**Figure 5.**
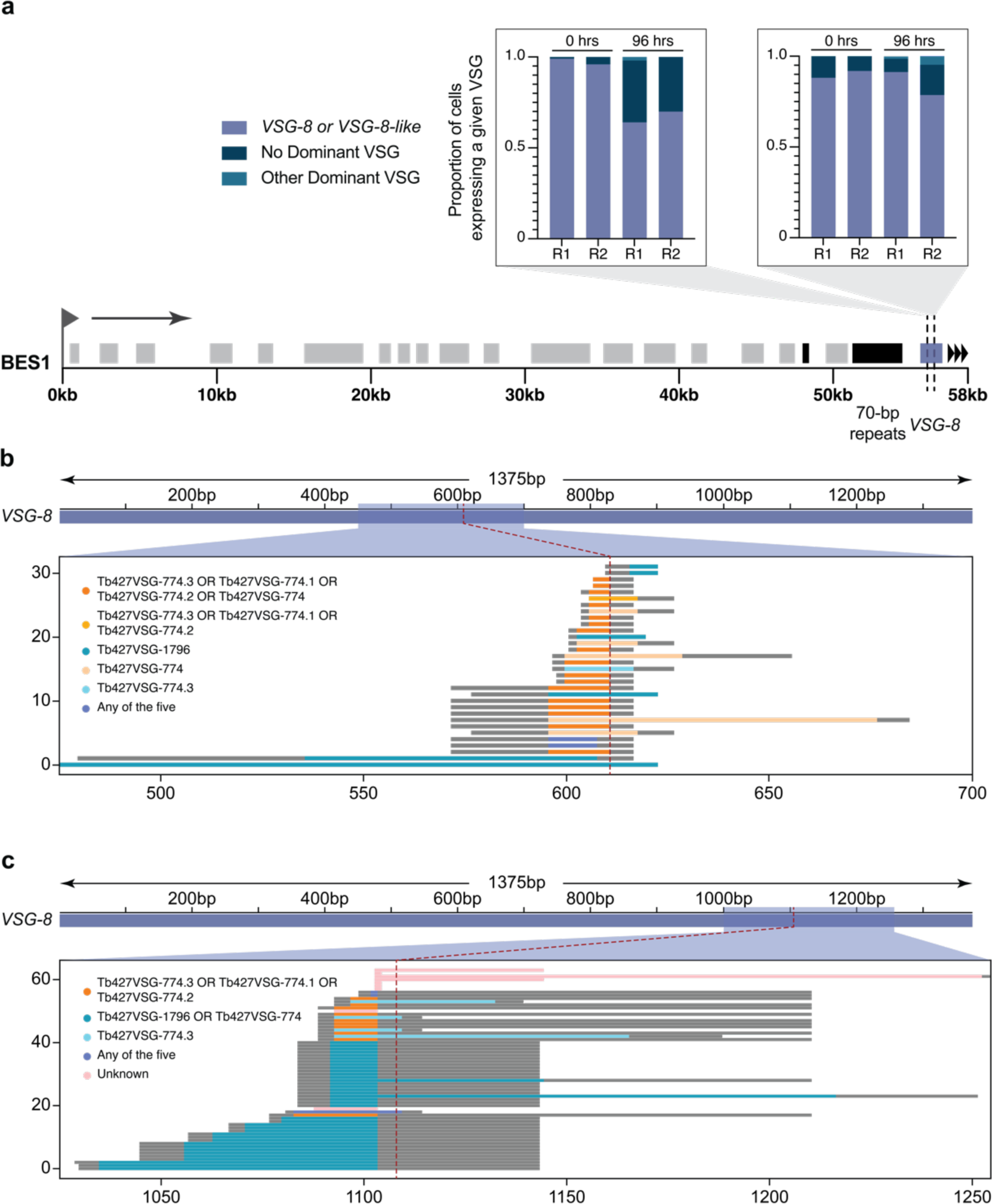
DSB in *VSG-8* CDS leads to switching by segmental gene conversion. **a,** Proportion of cells expressing a given VSG at 0 and 96 hours after DSB induction at nucleotide position 609 and 1105 of *VSG-8* CDS. The data is shown for two replicates per timepoint. **b, c** Recombinant fragments based on *de novo* assembled VSG transcripts for individual cells after 96 hours DSB induction at nucleotide positions 609 **(b)** and 1105 **(c)** of *VSG-8* CDS. The potential ‘donor’ VSG and the integrated stretch length are depicted by the colored lines. Gray extensions on the lines represent the maximal sequence length that could have been recombined (i.e. stretch until the next nucleotide difference between *VSG-8* and the possible donor(s)).

Analyzing the sequencing reads, we were surprised to find that >99.4% cells induced to have a DSB in the *VSG-8* CDS had retained BES1 transcription over its entire length, i.e. there was no evidence of DNA recombination within the 70-bp repeats nor of *in situ* switching, as we had observed for the *VSG-2* cut sites (Fig. 5a). Interestingly, however, the reads mapping onto the actively expressed VSG at 96 hours post-Cas9 induction presented a small number of SNPs around the expected DSB site when comparing them to the *VSG-8* sequence expressed in uninduced cells. Using the scRNA-seq reads, we generated *de novo* VSG assemblies for the individual cells and found that the location, identity, and number of SNPs varied between the sequenced cells, suggesting that the observed repair had occurred in a slightly different manner in each cell (Extended Data Fig. 5c,d). It is important to note that we never observed SNPs around the cut site when we induced a DSB in the *VSG-2* CDS. Since non-homologous end-joining does not occur in *T. brucei*^37^ and we observed no evidence of DNA resection and HR upstream of the break site, we investigated whether segmental gene conversion, using one of the (pseudo)genes similar to *VSG-8* as a template, was responsible for the observed generation of sequence variation at the active VSG gene.

Using BLAST, we searched for sequences similar to that of *VSG-8* and aligned them to the *de novo* assembled VSG transcripts that we had generated for each single cell. Interestingly, we observed that most of the novel sequence stretches around the DSB position had an exact match in at least one of the VSG genes similar to *VSG-8* present in the genome. The multiple sequence combinations observed suggests that a) the same VSG gene was not always used as the repair template and that b) the transferred DNA segment was of variable length (Fig. 5b,c). The VSGs that contained the matching SNPs were all pseudogenes and were located in different subtelomeric arrays within the genome. Thus, our data suggest that in the presence of a suitable repair template (VSG gene or pseudogene), DSBs in the active VSG gene are preferentially repaired by segmental gene conversion, leading to the generation of new ‘mosaic’ VSG genes.

To confirm that our observed switching phenotype was not unique to *VSG-8* expressing cells, we selected another switched clone expressing *VSG-11* in BES1 and designed sgRNAs to induce DSBs at nucleotide positions 519 or 729 of the *VSG-11* CDS. *VSG-11* also has a second copy in the genome and shares regions of DNA sequence homology with at least one other VSG. Sanger sequencing of VSG cDNA amplified from cells induced to generate breaks at either site on VSG-11 confirmed that mosaic VSGs had again been generated by segmental gene conversion (Extended Data Fig. 5d).

The observation that almost every cell expressing *VSG-8* or *VSG-11* repaired its DSB in the VSG CDS by segmental gene conversion, rather than by BIR as we observed for DSBs in *VSG-2*, suggests that *T. brucei* first searches for homologous VSG sequences to use as DNA repair templates before starting to resect the BES toward the 70-bp repeats.

## Discussion

To investigate the mechanism of VSG selection, we leveraged CRISPR-Cas9 technology to create targeted DSBs at specific sites along BES1 and developed a highly sensitive, trypanosome-tailored scRNA-seq protocol. This combination of techniques allowed us to uncover patterns of VSG expression and recombination before, during, and after a switch. Our findings indicate that the VSG selection mechanism is influenced by the DSB’s location relative to the 70-bp repeats and the presence (or absence) of DNA homology regions for repair.

From these results, we propose a model with two potential outcomes for a DSB in an active VSG gene: 1) If a homologous region exists in the genome, the DSB is repaired through segmental gene conversion, involving crossovers up- and downstream of the break site. This process often results in mosaic VSGs. 2) In the absence of a homologous region, the DSB is repaired via BIR, involving a single crossover in the 70-bp repeats. This leads to the duplication and activation of telomere-adjacent VSG genes from other telomeric locations (BESs, MESs, subtelomeric VSG arrays, or minichromosomes) into the active BES (see Fig 6).

**Figure 6.**
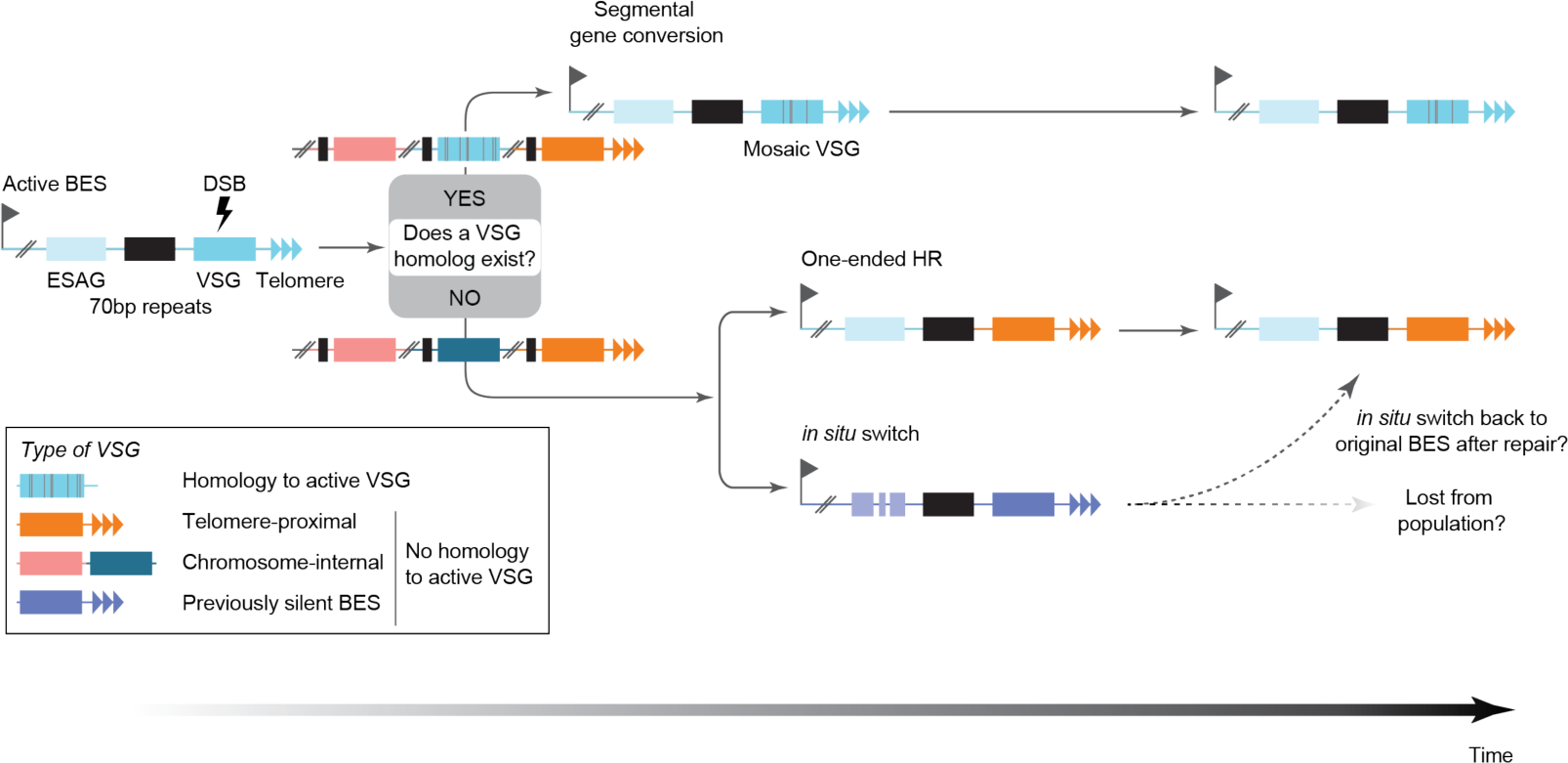
Model of VSG selection mechanism.

This model is supported by our observations following DSBs in actively transcribed *VSG-8* or *VSG-11* genes, for which homologous regions are available in different genomic loci. In these cases, segments from pseudogenes in subtelomeric VSG arrays were incorporated into the active VSG, forming mosaic VSGs. Interestingly, no VSGs from other BESs were copied via BIR, suggesting that *T. brucei* possesses an effective homology search mechanism and prefers repair by segmental gene conversion over BIR at the 70-bp repeats.

Our data also showed that DSBs in the actively transcribed *VSG-2*, which is atypical among VSGs in that it shares very little homology with other VSG genes, were repaired by BIR. Immediately after DSB induction, we observed a strong decrease in total VSG transcript levels and in BES1 ESAG transcripts, probably caused by a break-induced transcriptional arrest of the BES while the homology search was ongoing. At the same time, we observed a high proportion (>40%) of *in situ* switchers, i.e. cells that had activated transcription of another BES. Since we detected almost no *in situ* switchers at later time points (<3% at day 10), we speculate that they are either lost from the population due to a fitness disadvantage or that they represent an intermediate stage to ensure sufficient VSG transcript levels while BES1 is repaired and switch back to BES1 transcription once it is repaired. Previously, it was found that after inhibiting transcription in the active BES, it remains open (nucleosome-depleted), facilitating the ability of cells to reactivate it^38^. Either scenario would suggest that in Lister 427 isolates, transcription of BES1 confers a strong fitness advantage *in vitro* over transcription of other BES. This apparent selective pressure for BES1 may explain the low number of VSG switchers observed in this isolate compared to other isolates.

Combining CRISPR-Cas9 technology to induce cuts in both *VSG-2* and non-*VSG-2* expressing cell lines with transcriptional profiling of single cells, we were able to reveal the critical role of available homologous DNA sequence in determining the outcome of DSB repair. Furthermore, our findings suggest that mosaic formation might be the ‘preferred’ mechanism for antigen activation following a DSB in the active VSG. It is fast, does not involve resection and keeps the original BES active. Mosaic VSG formation is hypothesized to emerge as natural infections progress, serving as a mechanism for diversifying the range of antigenically distinct VSG isoforms^14,39^. While the exact mechanisms driving mosaic formation are still not completely understood, our results indicate that DNA homology plays a fundamental role in this process. It has been suggested that mosaic formation could occur either shortly before the functional VSG is recombined into the active BES, or within the active BES itself at the time of the VSG switch^14^. Our findings suggest the latter may be the case. Existing γ-H2A ChIP-seq and Breaks Labelling In Situ and Sequencing (BLISS) data indicate that DSBs naturally occur within the active VSG CDS^40,41^. Therefore, it seems plausible that mosaic formation occurs more frequently *in vitro* than previously recognized, particularly given the extensive research conducted on the atypical VSG isoform, VSG-2.

In summary, our work sheds light on the decades-old question of why some VSGs are more prevalent during infection than others. We found that the availability of homologous DNA for DSB repair and the genomic location of VSGs are key factors in determining the hierarchy of VSG activation. In addition, our study of the transcriptional changes immediately following a DSB in a VSG suggests that *in situ* switching may act as a temporary backup to maintain VSG transcript levels while the BES1 undergoes repair. Central to these discoveries was the development of the highly sensitive and precise SL-Smart-seq3xpress approach. This method enabled the measurement of VSG expression, whilst also revealing the intricate DSB repair process in single cells.

Importantly, our data also demonstrate that tailored, highly sensitive scRNA-seq approaches not only facilitate the study of cell-to-cell heterogeneity at the level of the transcriptome but to also discover underlying genomic changes. We believe that ability to link genomic and transcriptional diversity of pathogens at the single-cell level will be a powerful tool to dissect the evolution of drug resistance and to aid in the design of more robust drugs.

## Methods

### Trypanosome culture and genetic manipulation

Bloodstream form *T. brucei* Lister 427, 2T1^T7^ and 2T1^T7^ ^Cas9^ trypanosomes^42, 43^, along with their derivatives, were maintained at 37°C and 5% CO_2_ in HMI-11 supplemented with 10% FCS and the appropriate selective drug^44^. Cells were maintained below 2.0 × 10^6^ cells/ml. Cells were electroporated as previously described^45^. The 2T1^T7^ ^Cas9^ cell line was generated by transfecting 2T1^T7^ cells with the pRPa^Cas9^ plasmid^43^ which integrates into a tagged rDNA spacer region^42^. Cas9 expression was induced with 1µg/ml doxycycline. For the SL-Smart-seq3xpress switching assays, Cas9 induction was maintained throughout the time course. To generate clonal populations of VSG switched cells following the induction of a DSB, the induced population was diluted to 8-10 cells/ml in HMI-11 and spread across the wells of a 96-well plate in 100µl volumes. Plates were left for 5 days for the cells to recover. Cells were counted with a Beckmann Coulter cell counter.

### Sanger sequencing of expressed VSG transcripts

RNA was extracted from 5.0 × 10^6^ cells using the NucleoSpin RNA Kit (Macherey & Nagel) according to the manufacturer’s instructions. RNA was stored at −80°C. First strand cDNA was synthesized from 5µg of the extracted RNA using the SuperScript II Reverse Transcriptase (Invitrogen) and the oligo(dT)_12-18_ primer as per the manufacturer’s instructions. First strand cDNA was stored at −20°C. Expressed VSG transcripts were amplified from the first strand cDNA by PCR using a forward primer specific to the spliced leader sequence (5’-GACTAGTTTCTGTACTAT-3’) and a reverse primer specific to the conserved 3’ sequence of VSG mRNAs (5’-CCGGGTACCGTGTTAAAATATATC-3’). For each PCR reaction, 1µl of 1:50 diluted first strand cDNA was used. PCR products were visualized following agarose gel electrophoresis and the VSG amplicon purified from the agarose using the Nucleospin Gel and PCR Clean-up Kit (Macherey & Nagel) as per the manufacturer’s instructions. Sanger sequencing of at least 100ng of purified PCR product was performed by Eurofins Genomics using either of the primers used in the VSG amplification PCR.

### sgRNA design and cloning

sgRNA target sequences were designed with Protospacer Workbench^46^ using the Lister 427 2018 genome assembly (^10^, https://tritrypdb.org/) as the reference database and optimized for use with SpCas9. sgRNA target sequences were selected according to their Bowtie Score (a measure of off-target Cas9 activity) and the Doench-Root-Activity score (a measure of sgRNA activity). As described in Rico et al^43^, ‘aggg’ was added to the forward primer sequence and ‘caaa’ to the reverse primer sequence to create the BbsI cloning sites. The target sequences were cloned into the pT7^sgRNA^ plasmid and transfected into the 2T1^T7^ ^Cas9^ cell line as previously described^43^. The pT7^sgRNA^ plasmid integrates into a random rDNA spacer region. Since up to 20 of these regions exist in the genome, this allows for the transfection of the same cell line with multiple different sgRNAs simultaneously.

### Western blot analysis of Cas9 and γ-H2A expression

Total protein extract from 2.0 × 10^6^ cells was boiled in 1X lysis buffer (1:3 4X Laemmli:1X RIPA, 2mM DTT, 1% β-mercaptoethanol) and separated on a 12.5% SDS-PAGE gel. Separated proteins were transferred onto a methanol-equilibrated PVDF membrane using a Bio-Rad Mini Trans Blot Cell according to the manufacturers’ instructions. To visualize transferred proteins, the membrane was stained with 0.5% Amido black solution (in 10% acetic acid). Destaining was performed with 1X destaining solution (25% isopropanol, 10% acetic acid). The blotted PVDF membrane was cut into three according to the prestained protein ladder before blocking: above 70kDa for the detection of SpCas9, between 70kDa and 25kDa for the detection of the EF1α loading control and below 25kDa for the detection of γ-H2A. Cas9 and EF1-α blots were blocked in 5% milk/PBS-T and γ-H2A blots were blocked in 3% BSA/PBS-T at room temperature. The membranes were washed three times with 1X PBS-T and primary antibody incubation performed overnight at 4°C. The primary antibodies were used at the following dilutions: anti-Cas9 (1:1000 in 5% milk/PBS-T, Active Motif, clone 7A9-3A3); anti-EF1α (1:20,000 in 1% milk/PBS-T, EMD Millipore Corporation, clone CBP-KK1); anti-γ-H2A (1:200 in 1% milk/PBS-T). After washing the membranes 3 more times with PBS-T, the following HRP conjugated secondary antibodies were used: for Cas9 - anti-mouse (1:10,000 in 1% milk PBS-T, GE Healthcare, code NA931V); for γ-H2A - anti-rabbit (1:2000 in 1% milk/PBS-T, GE Healthcare, code NA934V); for EF1-α - anti-mouse (1:10,000 in 1% milk/PBS-T, GE Healthcare, code NA931V). Following secondary incubation, the membrane was washed three times with PBS-T and once more with PBS. For signal detection, the Immobilon Western chemiluminescent HRP substrate was used according to manufacturers’ instructions. The signal was visualized on a ChemiDoc MP Imaging System.

### Immunofluorescence analysis of γ-H2A expression

Immunofluorescence analysis of γ-H2A expression was performed as previously reported^47^. At least 250 cells were analyzed per sample. Images were acquired with a Leica DMi8 inverted fluorescence microscope and processed with Fiji software.

### FACS analysis of VSG expression

FACS analysis of VSG-2 expression was performed on live cells and therefore all steps were performed at 4°C to prevent internalization of the VSG-2 antibody. Cells were stained immediately prior to analysis. For each replicate, 1.0 × 10^6^ cells were harvested by centrifugation and incubated with fluorescently-conjugated anti-VSG-2^48^ diluted 1:500 in HMI-11 in the dark. Cells were washed three times with 1X TDB and resuspended in 400µl of 1X TDB. Cells were stained with 1µg/ml propidium iodide for the identification of dead cells. Samples were processed on a FACS Canto (BD Biosciences) and 10,000 events were captured per sample. Data was processed using FCS Express software. Gates were applied to remove cellular debris (FSC-A vs SSC-A) and remove doublets (FSC-A vs FSC-H).

### Single cell sorting for SL-Smart-seq3xpress library preparation

Single cells were sorted into 384-well plates for SL-Smart-seq3xpress library preparation by flow cytometry using a FACS Fusion II cell sorter (BD Biosciences) and a 100µm nozzle within a safety cabinet. The sorter was calibrated according to the manufacturer’s protocol prior to harvesting cells to reduce the time cells were held prior to sorting, thereby reducing cell death. A 384-well plate adapter was installed and pre-chilled to 4°C. Correct droplet positioning within wells was verified visually by sorting empty droplets onto a covered 384-well plate before every plate was sorted.

5.0 × 10^6^ cells were harvested by centrifugation at 4°C and washed twice in sterile filtered ice cold 1X TDB. The cells were resuspended in 1ml of ice cold filtered 1X TDB, stained with 1µg/ml propidium iodide and brought immediately to the sorter on ice. Populations were gated to remove cellular debris, doublets and dead cells as described above. As a consequence of our tight gating strategy (Extended Data Fig. 6), we likely enriched for cells in G1 and excluded larger cells in G2. Plates prepared with lysis buffer were thawed individually immediately prior to sorting and placed within the precooled adapter. Single cells were sorted using the ‘single cell’ purity option into the appropriate wells and the plates were immediately sealed with an aluminum foil and moved to dry ice before longer term storage at −80°C. Sorted plates were not stored for more than one month prior to library preparation.

### Generation of SL-Smart-seq3xpress RNA spike-ins

From the standard ERCC RNA spike-in set of 92 sequences, 10 with a size of around ∼500 nt and ∼50% ∼G+C content were selected and synthetic spike-in DNA fragments were ordered from IDT, adding a homology region for cloning, the T7 promoter sequence and the SL sequence on the 5’ end and a homology region for cloning on the 3’ end. The fragments were cloned into a pBSIIKS+ plasmid digested with SacI and BamHI (NEB) using Infusion (Takara) and transformed into Stellar cells. Plasmids were then extracted, linearized with BamHI, and *in vitro* transcription and polyadenylation was performed using HiScribe T7 ARCA mRNA Kit (with tailing) from NEB, following recommended procedure. The obtained RNA from each of the 10 spike-in sequences were mixed and aliquot dilutions of the spike-in mix were generated and stored at −80°C until usage.

### SL-Smart-seq3xpress library preparation

RNase free reagents were used for all steps and all surfaces were regularly treated with RNaseZAP (Sigma). Each well of a 384-well plates was filled with 3µl/well silicone oil (Sigma) using an Integra Assist Plus pipetting robot. The plates were sealed with adhesive PCR plate seals and briefly centrifuged to collect the liquid at the bottom of the wells. To each well, 0.3µl of lysis buffer (0.1% TX-100, 5% w/v PEG8000 adjusted to RT volume, 0.008µM oligodT (5’-Biotin-AGAGACAGATTGCGCAATG[N_8_][T_30_]VN-3’) adjusted to RT volume, 0.5mM each dNTP adjusted to RT volume, 0.5U/µl RNase inhibitor, 0.033µl of spike-in mix (∼1364 transcripts)) was added using an I.DOT liquid dispenser (Cytena). The plates were briefly centrifuged, placed on ice and brought to the cell sorter. Single cells were sorted into each well as described above.

To lyse cells, the reaction plate was thawed, centrifuged, incubated at 72°C for 10 minutes. To each well, 0.1µl of reverse transcription mix (25mM Tris-HCl pH8.3, 30mM NaCl, 2.5mM MgCl2, 8mM DTT, 0.25U/µl RNase inhibitor, 2U/µl Maxima H-minus RT) was immediately added following cell lysis using the I.DOT and the reaction plate then incubated at 42°C for 90 minutes before inactivation of the reaction at 85°C for 5 minutes. The second strand cDNA synthesis and PCR amplification was performed using primers that annealed to the SL sequence (SL primer, 5’- Biotin-ACACTCTTTCCCTACACGACGC-3’) and a conserved sequence added by the oligodT primer (5’- GTCTCGTGGGCTCGGAGATGTGTATAAGAGACAGATTGCGCAATG-3’).

Immediately following the reverse transcription reaction, the reaction plate was centrifuged and 0.6µl of PCR amplification mix (1X SeqAmp PCR buffer, 0.025U/µl SeqAmp polymerase, 0.5µM SL primer, 0.5µM Reverse primer) added to each well using the I.DOT liquid dispenser. The PCR reaction was performed with the following conditions: 95°C for 1 minute, 16 cycles of (98°C for 10 seconds, 65°C for 30 seconds, 68°C for 4 minutes), 72°C for 10 minutes). Following the reaction, the PCR plate was centrifuged and amplified cDNA was then diluted by adding 9µl of dH_2_O to each well of the plate. If not used immediately, the plate was stored at −20^°^C until next step.

To each well of a fresh 384-well plate, 1µl of each prediluted cDNA was added using the Integra pipetting robot. To each well, 1µl of tagmentation mix (10mM Tris-HCl pH7.5, 5mM MgCl2, 5% DMF, 0.002µl TDE1) was added and the plate incubated at 55°C for 10 minutes. To stop the reaction, 0.5µl of 0.2% SDS was immediately added to each well. The plate was centrifuged and incubated at RT for 5 minutes. The individual libraries generated from each well were dual indexed with Illumina i5 (5’- AATGATACGGCGACCACCGAGATCTACAC[8bp index]TCGTCGGCAGCGTC −3’) and i7 index primers. For each index primer, 0.5µl (2.2µM) was dispensed into each of the reaction wells and 1.5µl of PCR mix (1X Phusion HF Buffer, 0.2mM each dNTP, Tween-20 0.025% and 0.01U/µl Phusion HF DNA polymerase, final concentrations in the 5µl reaction) added to each of the wells. The PCR reaction was performed with the following conditions: 72°C for 3 minutes, 95 degrees for 30 seconds, 14 cycles of (95°C for 10 seconds, 55°C for 30 seconds, 72°C for 1 minute), 72°C for 5 minutes). The reaction volume from all wells were pooled into a robotic reservoir (Nalgene, Thermo Scientific) by centrifuging the plate placed in custom-made 3D printed plate holder at 200g for 30 seconds. The pooled library was purified using AMPure XP beads at a ratio of 1:0.7. The libraries were eluted from the beads in 45µl total volume of dH_2_0. To further decrease free unligated adapter concentration, the libraries were run on a 4% non-denaturing PAGE gel and purified according to standard polyacrylamide gel purification protocols. The libraries were sequenced on a NextSeq 1000 sequencing platform to produce paired-end reads of 101nt (cDNA read) and 19nt (TAG+UMI read), and 8nt for the index reads.

### 5’ Chromium 10X library preparation and sequencing

Cultures of N50 and P10 cells were set up and maintained at 0.5-1.0 × 10^6^ cells/ml prior to harvesting for library preparation. A mixed population sample was prepared by pooling together equal numbers of N50 and P10 cells. The mixed cells were harvested by centrifugation at 400g for 10 minutes, washed twice in ice-cold 1X PBS supplemented with 1% D-glucose and 0.04% BSA, and resuspended in 1ml of the buffer. The cells were then filtered with a 35µm cell strainer (Corning) and adjusted to 1000 cells/µl. Libraries were prepared using the Next GEM Single Cell 5’ GEM Kit v2 (10xGenomics) and sequenced on the NextSeq 1000 platform to a depth of ∼50,000 reads per cell. Paired-end reads of 26nt (read 1) and 122nt (read 2) as well as 10nt-index reads were generated.

### Primary processing of SL-Smart-seq3xpress sequencing data

The two reads containing the indexes (8nt each) and the TAG+UMI(19nt) were concatenated into a 35nt read. Artefact reads containing the TAG sequence (or its reverse complement) in the cDNA read were filtered out using Cutadapt^49^. Downsampling of reads for method benchmarking was done using seqtk^50^. Reads were mapped with STARsolo^51,52^ (STAR version 2.7.10a) to a hybrid fasta file combining the *T. brucei* Lister 427 strain genome (Tb427v11^5^) and the spike-in sequences, producing a transcript count matrix and an alignment (BAM) file. The count matrix was then corrected using the index hopping filtering pipeline “scSwitchFilter” (described in the next section) using the BAM file as input.

### Index hopping filtering

scSwitchFilter corrects index hopping in multiplexed sequencing libraries using raw BAM files instead of a count matrix^53^. The correction process involves three major steps: 1) BAM to SAM conversion; 2) read extraction and parsing; 3) negative correction count matrix computation. In the first step the pipeline utilizes samtools to convert a BAM file to a SAM file. In step 2, a fast bash script is employed to extract and parse valid reads from the SAM file, select reads with CB (cell barcode), UB (UMI barcode) and GN (gene name) tags, and split CB tags for subsequent analysis. The selected reads are then complied into a single TSV file. Depending on the number of plates (individual libraries) in the sequencing experiment (run), the script may split the CB tag into plate/library-i5-i7 or i5-i7 barcode combinations. In step 3, scSwitchFilter calculates read counts for switched indices, assuming a low probability of the combination of UB, GN, and an index being present in multiple plate wells. Reads with over 80% (default threshold) of total counts among those with switched indices remain unfiltered. The tool generates a residue count matrix that should be subtracted from the initial count matrix to obtain the filtered count matrix.

### SL-Smart-seq3xpress data analysis

Count matrices were processed with JupyterLab (version 3.4.4) notebooks using IPython (version 7.31). Cells with less than 500 genes detected, 1000 gene UMI transcript counts and 50 spike-in UMI counts were filtered-out. For the gene expression analysis, transcript counts for each cell were normalized by spike-in counts. For the quantification of cells expressing each VSG, we defined a cell as expressing a given VSG, if the transcript counts for that VSG represented more than 80% of the transcript counts for all VSGs in that cell. If no VSG reached this threshold, we defined the cell as having ‘no dominant VSG’.

### Sensitivity and specificity comparison for different sequencing approaches

For Smart-seq2 and SL-Smart-seq3xpress data, reads were subsampled to match the average reads per cell in Chromium 10X (∼75K reads/cell in Briggs et al^28^ and ∼100K reads/cell for Chromium 10X data from this study). All sequencing data was mapped with STARsolo with identical settings. For the sensitivity comparison, the transcript end coordinate annotations were extended until the beginning of the next transcript, matching the conditions used by Briggs et al. For specificity comparison, cells with either genes detected, total transcript counts or total VSG transcript counts below half of the median for the population of cells, were filtered out. Additionally, only cells with >10 VSG UMI counts were considered.

### Type of switching analysis

Single-cell BAM files were extracted from STARsolo output BAM file, using the cell-specific CB:Z attribute (storing the indexes and the TAG sequence) for each mapped read in the BAM file. Only cells with a dominant VSG from 0hs, 96hs and 10days post-induction time points were considered. Coverage files (Bigwig files) were generated for each single-cell using deepTools^54^ (version 3.5.4) bamCoverage function with “-- normalizeUsing RPKM” and “--minMappingQuality 10” options. Coverage tracks were plotted using pyGenomeTracks^55^. For the determination of switching type (recombination or *in situ*) and for the identification of the transcriptional signal end position in BES1, and the start of transcriptional signal in the BES where the newly active VSG was originally located, the single-cell coverage tracks were visually inspected in IGV^56^ ( version 2.16.0).

### Identification of VSG homologs

Homologs of *VSG-2*, *VSG-8* and *VSG-11* were identified by BLAST to the Lister 427 genome assembly in TriTrypDB^57^. Hits with a bitscore > 1000 were selected as highly similar homologs and putative ‘donors’ for segmental gene conversion. For *VSG-*2, *VSG-8* and *VSG-11*, there were 0 hits, 5 hits and 1 hit meeting this criterion, respectively.

### Single-cell *de novo* VSG transcript assembly after DSB induction in *VSG-8*

Fastq files were demultiplexed into single-cell fastq files with deML^58^ (version 1.1.13) with default settings. De novo transcript assemblies were then generated for each single-cell with Trinity^59^ (version 2.15.1), restricting the output to contigs bigger than 1kb. To identify which of the assembled transcripts was the active VSG, the de novo assembled contigs were aligned with BLAST^60^ (version 2.14.0) to *VSG-8*, and the contigs with high similarity (bitscore > 2000) were selected. Those cells with no contig reaching the threshold were discarded. Multifasta files with all the single-cell *de novo* assembled VSGs per experiment, together with the putative ‘donor’ VSGs, were constructed, and aligned to VSG-8 with minimap2^61^ (version 2.10). Finally, the alignments were visualized in IGV and the start and end position of recombination and the putative donor(s) for each cell was determined.

### Bulk RNA-seq library preparation and sequencing

Cell lines expressing different VSGs – *VSG-2*, *VSG-8*, *VSG-11* – were maintained at 0.5-1.0×10^6^ cells/ml prior to harvesting. RNA-seq library preparation was performed as previously described^61^. Strand-specific RNA-seq library concentrations were measured in duplicate using Qubit dsDNA HS Assay Kit and Agilent TapeStation system. The libraries were quantified with the KAPA Library Quantification Kit according to the manufacturer’s protocol and sequenced on the Illumina NextSeq 1000 platform to generate paired-end reads.

### Bulk RNA-seq data analysis

Reads were mapped to the Lister 427 genome assembly v11 with STAR. Coverage files were generated and plotted in the same way as for the scRNA-seq data (see Methods section “Type of switching analysis“).

## Data availability

The scRNA-seq and RNA-seq data have been deposited in the European Nucleotide Archive and are accessible through ENA study accession number PRJEB72370.

## Code availability

Workflows, custom-made Unix Shell and Python scripts to analyse the data have been deposited into Zenodo (https://doi.org/10.5281/zenodo.10692101).

## Acknowledgments

We thank all members of the Siegel lab and the Division of Physiological Chemistry for valuable discussion. We thank the Laboratory for Functional Genome Analysis (LAFUGA) at the Gene Center Munich, LMU, for NGS, the Core Facility Flow Cytometry at the Biomedical Center, LMU, for providing equipment and expertise, and the BMC Core Facility Bioinformatics for providing access to the computing server. We thank Antoine-Emmanuel Saliba (Institute for RNA-based Research, Würzburg Germany) and Michael Hagemann-Jensen from the Sandberg Lab (Karolinska Institutet, Stockholm, Sweden) for advice on the adaption of the Smart-seq3xpress protocol. We thank Lucy Glover (Institut Pasteur, Paris, France) for providing the γ- H2A antibody and Eduardo Beltran (BMC, LMU) for 10x Genomics sequencing. This work was funded by the German Research Foundation [SI 1610 / 2-2], and an ERC Starting Grant (3D_Tryps 715466) and an ERC Consolidator Grant (SwitchDecoding 101044320) awarded to TNS. ZK and AD were supported by MSCA ITN Cell2Cell fellowships. KRM was supported by the European Union’s Framework Programme for Research and Innovation Horizon 2020 (2014-2020) under the Marie Skłodowska-Curie Grant Agreement No. 754388 (LMUResearchFellows) and from LMUexcellent, funded by the Federal Ministry of Education and Research (BMBF) and the Free State of Bavaria under the Excellence Strategy of the German Federal Government and the Länder.

## Author contributions

The experiments were designed by KRM, ZK, ROC, IS, MC-T and TNS and carried out by KRM, ZK, ROC and IS, unless otherwise indicated. Cell lines were generated by KRM and IS. The SL-Smart-seq3xpress approach was developed by KRM, ZK, ROC and TNS. SL-Smart-seq3xpress libraries were generated by ZK and ROC. Computational analyses were performed by ROC and AD. All authors contributed to the data interpretation and development of a model. The work was supervised by MC-T and TNS. The manuscript was written by KRM and TNS with help from ZK and ROC and edited by all other co-authors. The figures were generated by KRM, ZK and ROC.

## Additional information

Correspondence and requests for materials should be addressed to T. Nicolai Siegel.

## Extended Data Figures

**Extended Data Figure. 1.**
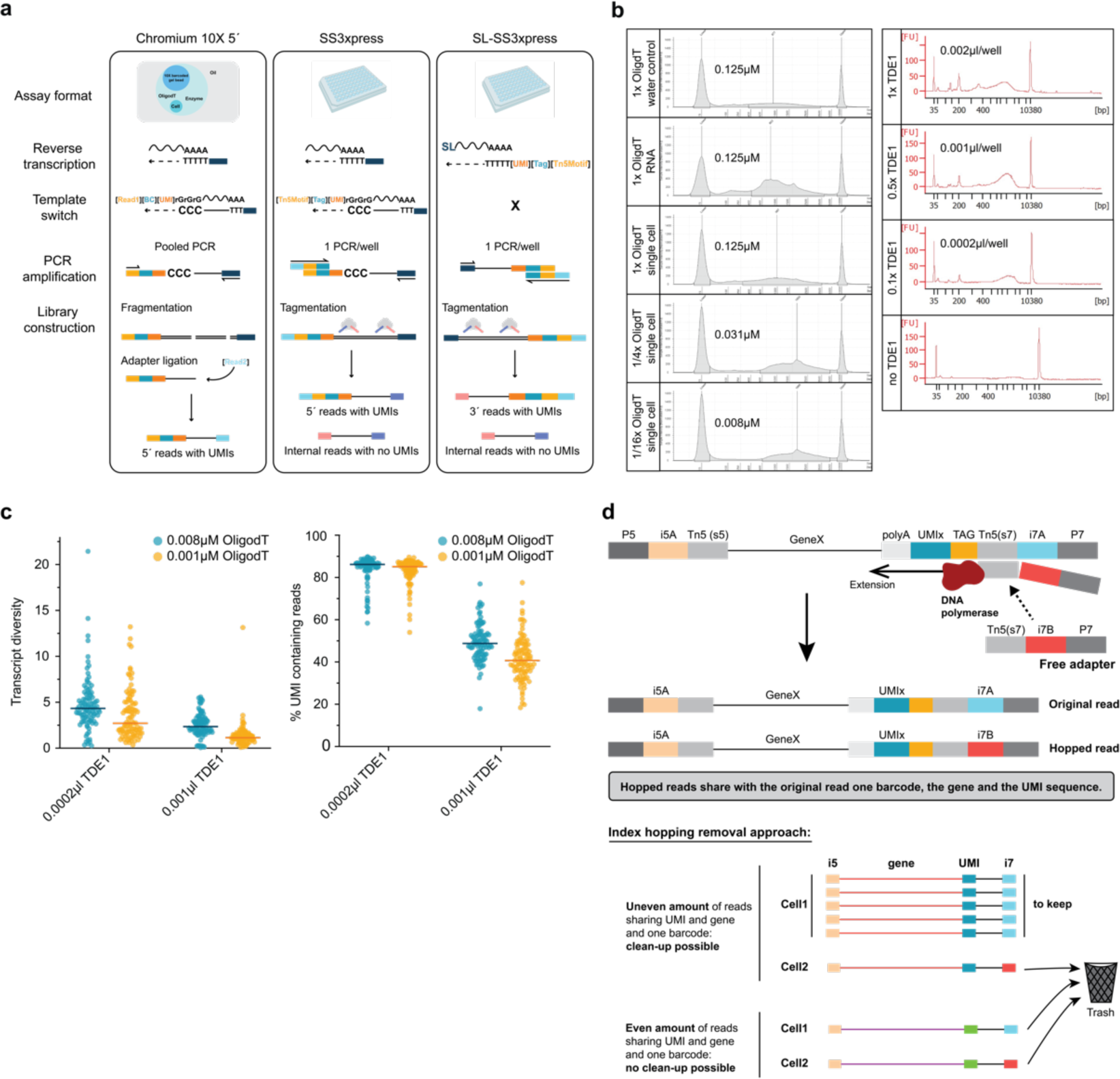
Optimization of SL-Smart-seq3xpress scRNA-seq pipeline. **a.** The library preparation workflows for: the Chromium 10X 5’ pipeline (left panel); the standard Smart-seq3xpress (SS3xpress) pipeline (middle); and the tailored SL-Smart-seq3xpress (SL-SS3xpress) pipeline (right). BC, barcode; SL, spliced leader. **b,** Left panel: Representative TapeStation profiles of single cell libraries (pooled and bead-purified after cDNA dilution) prepared using different oligodT concentrations - 1x, 1/4x, 1/16x - relative to the concentration in the published Smart-seq3xpress protocol^20^. Each condition was tested with at least two independent replicates, each containing between 6 and 48 cells. Right panel: TDE1 Tn5 concentrations −1x, 0.5x, 0.1x, relative to the concentration in the published Smart-seq3xpress protocol^20^ - tested to optimize tagmentation. Here, each condition contains 48 cells. **c,** Optimisation of SL-Smart-Seq3xpress oligodT and TDE1 enzyme concentration. Left panel: Transcript diversity of SL-Smart-Seq3xpress libraries prepared with varying oligodT and TDE1 concentrations. Transcript diversity was measured as % of UMI counts/per reads sequenced/single cell. Right panel: Percent of UMI containing reads in SL-Smart-seq3xpress libraries prepared with varying oligodT and TDE1 concentrations. Each dot represents a single cell. 96 cells were sequenced for each tested condition. **d,** A schematic of the mechanism leading to index hopping and our bioinformatic strategy to remove index hopped reads.

**Extended Data Figure 2.**
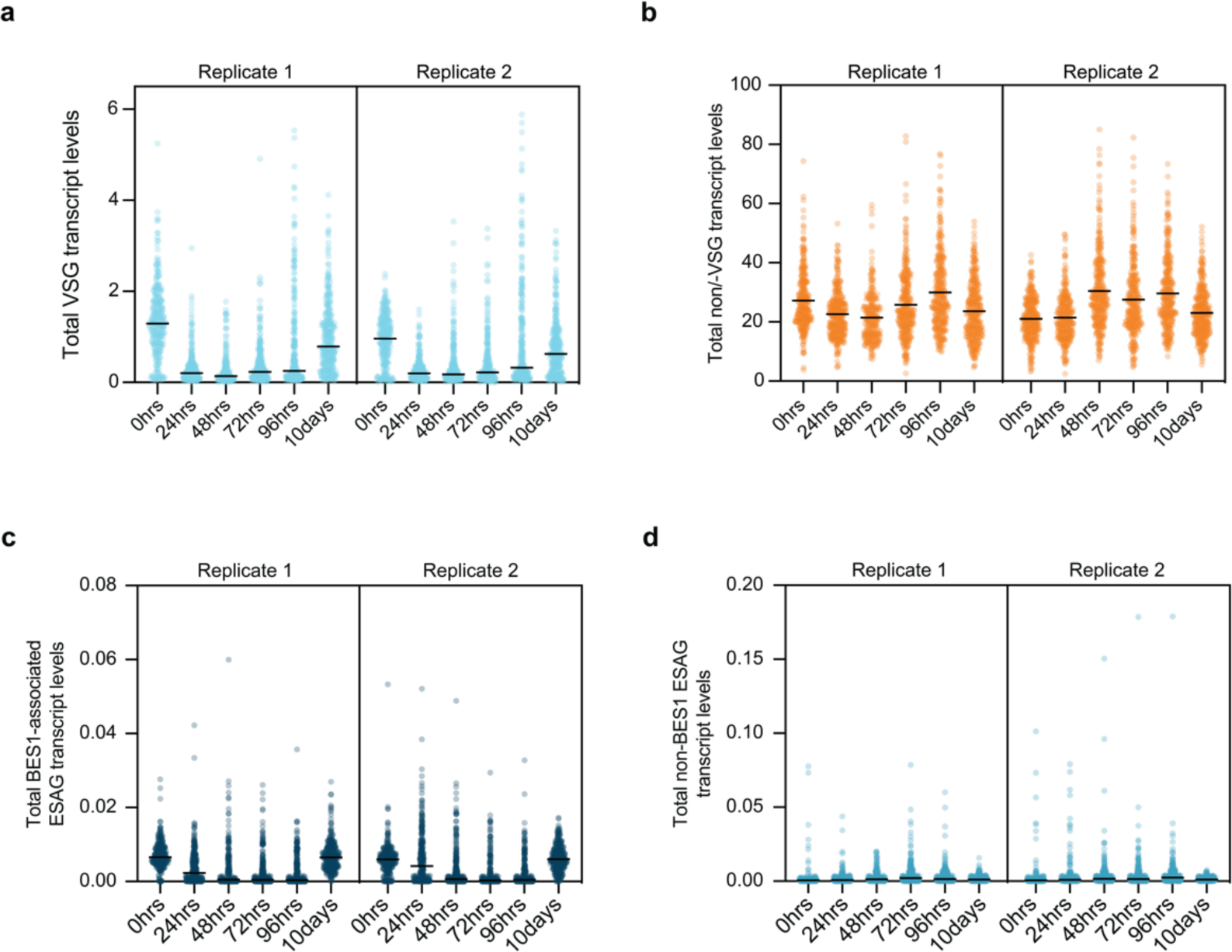
Relative expression levels of VSGs and ESAGs during the DSB induction in nucleotide position 152 from *VSG-2* CDS. **a-d,** Single cell transcripts level normalized by spike-in counts. The data is shown separately for each of two replicates (replicate 1 – R1, replicate 2 - R2). **a,** Total VSG genes, **b,** Total Non-VSG genes, **c,** Total BES1 ESAGs and **d,** Total Non-BES1 ESAGs.

**Extended Data Figure 3.**
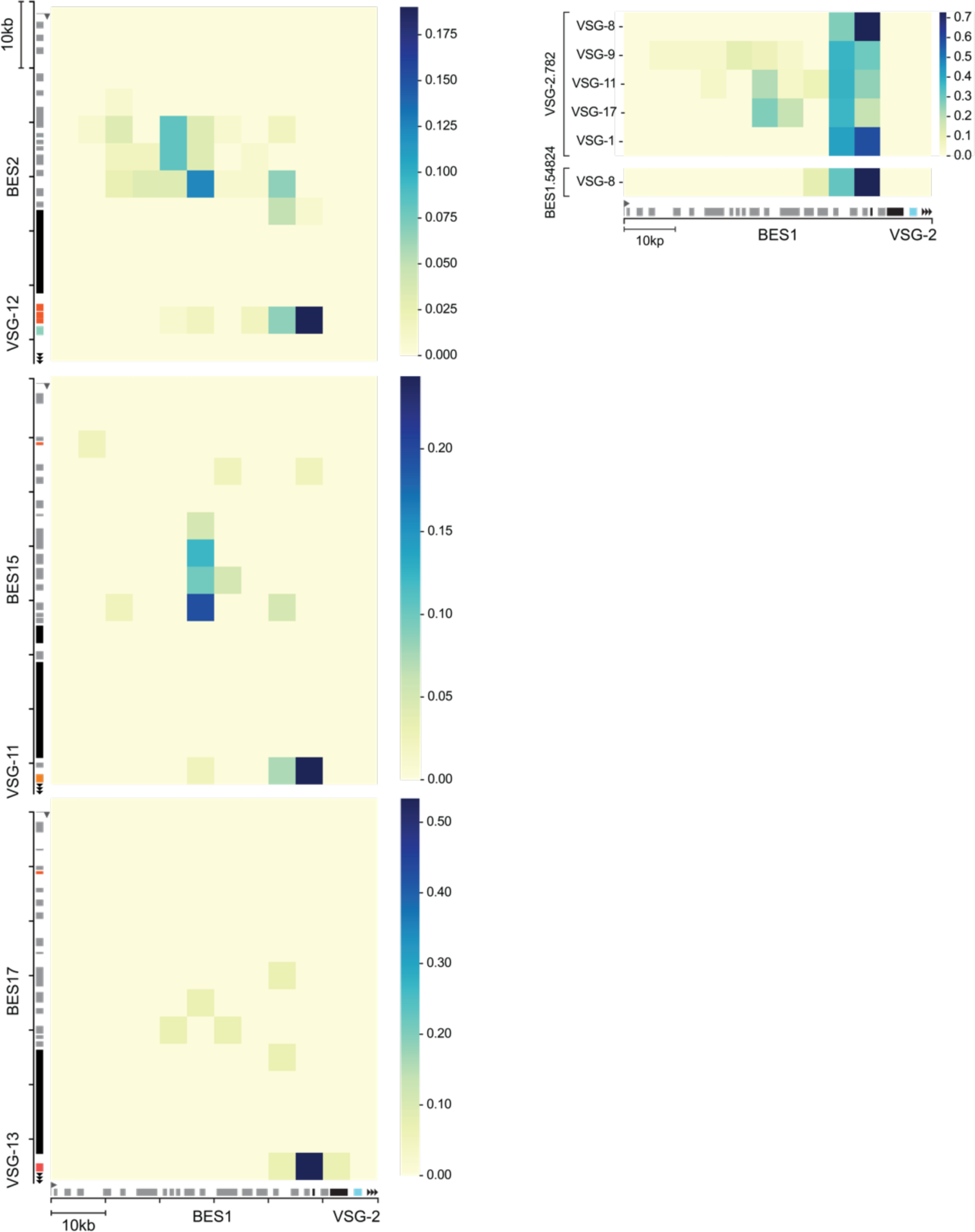
Summary of transcriptional signal end position in BES1 versus transcriptional signal start position in BES of incoming VSG for cells switching by recombination. **Left panel,** Heatmaps summarizing transcriptional signal end position on BES1 (columns) for recombinant switcher cells versus the transcriptional signal start on the BES from the incoming VSG, after a DSB in the active *VSG-2* CDS (at nucleotide position 152), as a proxy of recombination site preference. Each square on the heatmap represent 5kb bins (each column representing 5kb bins of BES1 and rows - 5kb bins of the BES from the incoming VSG). The color in each square represents the fraction of cells for which transcriptional signal ended at that given bin in BES1 and started at that given bin in the BES of the incoming VSG. The heatmaps summarize the data from 122 cells switching to *VSG-9*, 41 cells switching to *VSG-11* and 15 cells switching to *VSG-13*, at 96 hours and 10 days timepoints after DSB induction. **Right panel**, Heatmap summarizing transcriptional signal end position on BES1 for recombinant switcher cells after a DSB in nucleotide position 782 of *VSG- 2* CDS (VSG2.782) and nucleotide position 54824 of BES1 (BES1.54824). Each box in the heatmap represents the fraction of cells, for switchers to a given VSG (rows), which transcriptional signal ends at the given 5Kb bin of BES1 (columns). All VSGs expressed in at least 10 cells were considered.

**Extended Data Figure 4.**
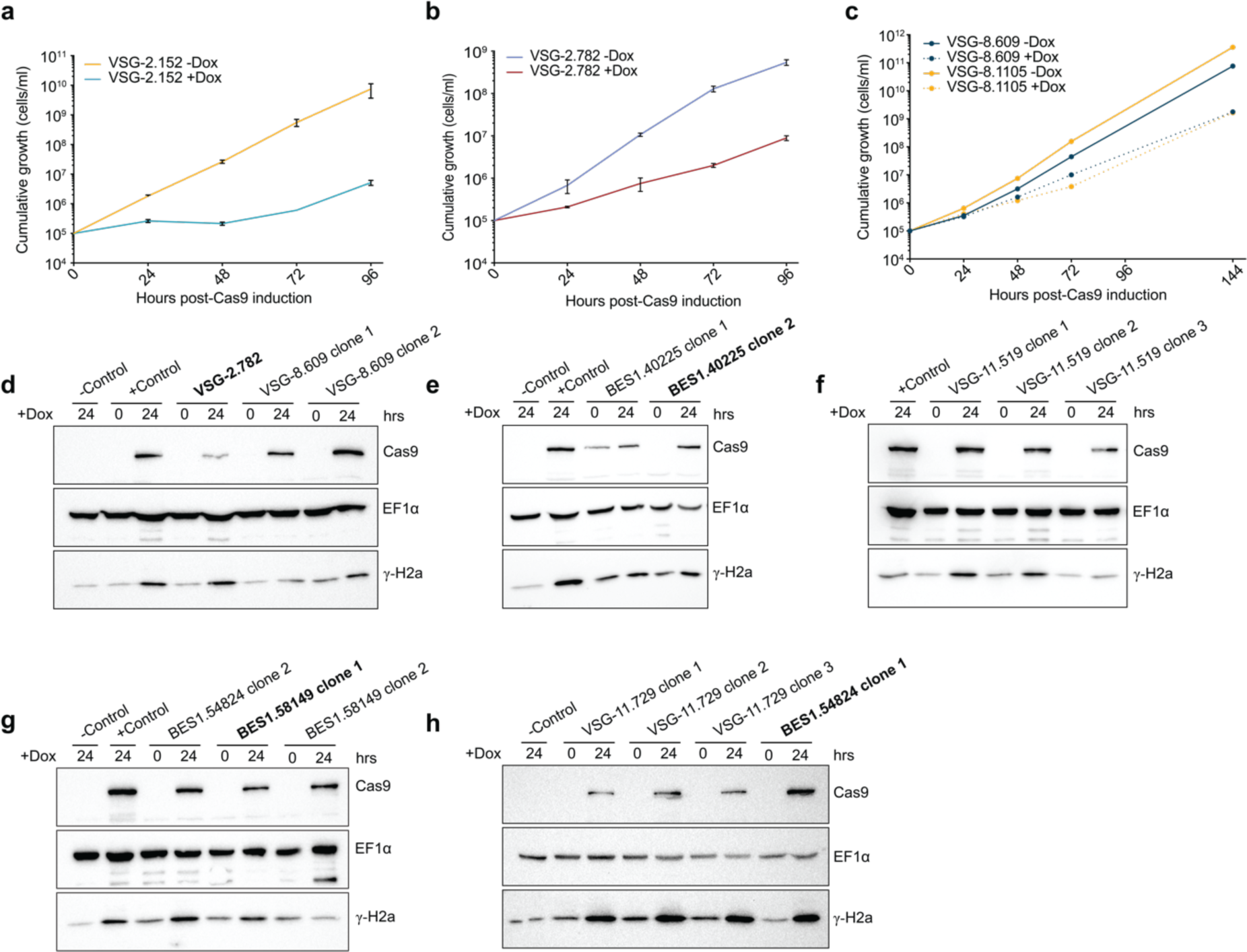
Growth curves and western blotting results after DSB inductions. **a-c,** Growth curves following Cas9-based DSB induction for cut sites at nucleotide position 152 (a) and 782 (b) of *VSG-2* CDS, and for cut sites at nucleotide position 609 and 1105 of *VSG-8* CDS (c). The values shown represent the mean of three biological replicates for VSG-2.152 and VSG-2.782 cell lines. For VSG-8 cell lines one replicate was tested. **e,** Heatmap summarizing transcriptional signal end position on BES1 for recombinant switcher cells after a DSB in nucleotide position 782 of *VSG-2* CDS (VSG-2.782) and nucleotide position 54824 of BES1 (BES1.54824). Each box in the heatmap represents the fraction of cells, for switchers to a given VSG (rows), which transcriptional signal ends at the given 5kb bin of BES1 (columns). All VSGs expressed in at least 10 cells were considered. **d-h,** Western blotting showing Cas9 and γ-H2a expression levels before and 24 hours post-Cas9 induction with doxycycline. Wild-type Lister 427 at 24 hours after doxycycline addition were used as a negative control. EF1α was used as a loading control. Clones used for scRNA-seq experiments are highlighted in bold. **d,** The results are shown for cut positions 152 and 782 in *VSG- 2* CDS and two independent clones for cut position 609 in *VSG-8* CDS. **e,** Cas9 and γ-H2a expression profiles for cut position 40225 (clones 1 and 2) in BES1 of a VSG-2 expressing cell line. **f,** Cas9 and γ-H2a expression levels for three clones of *VSG-11* expressing cell line with cut position at 519. For Sanger sequencing analysis VSG-11.519 clone 1 was used. **g,** The blot shows the results for *VSG-2* expressing cell lines with cut position at 54824 (clone 2) and 58149 (clones 1 and 2). **h,** The data is shown for *VSG-11* expressing cell line with cut position at 729 (clones 1, 2, and 3) and *VSG-2* expressing cell line with cut position at 54824 (clone 1). For Sanger sequencing VSG-11.729 clone 1 was used. Nucleotide positions are considered relative to the promoter in BES1 for cut sites

**Extended Data Figure 5.**
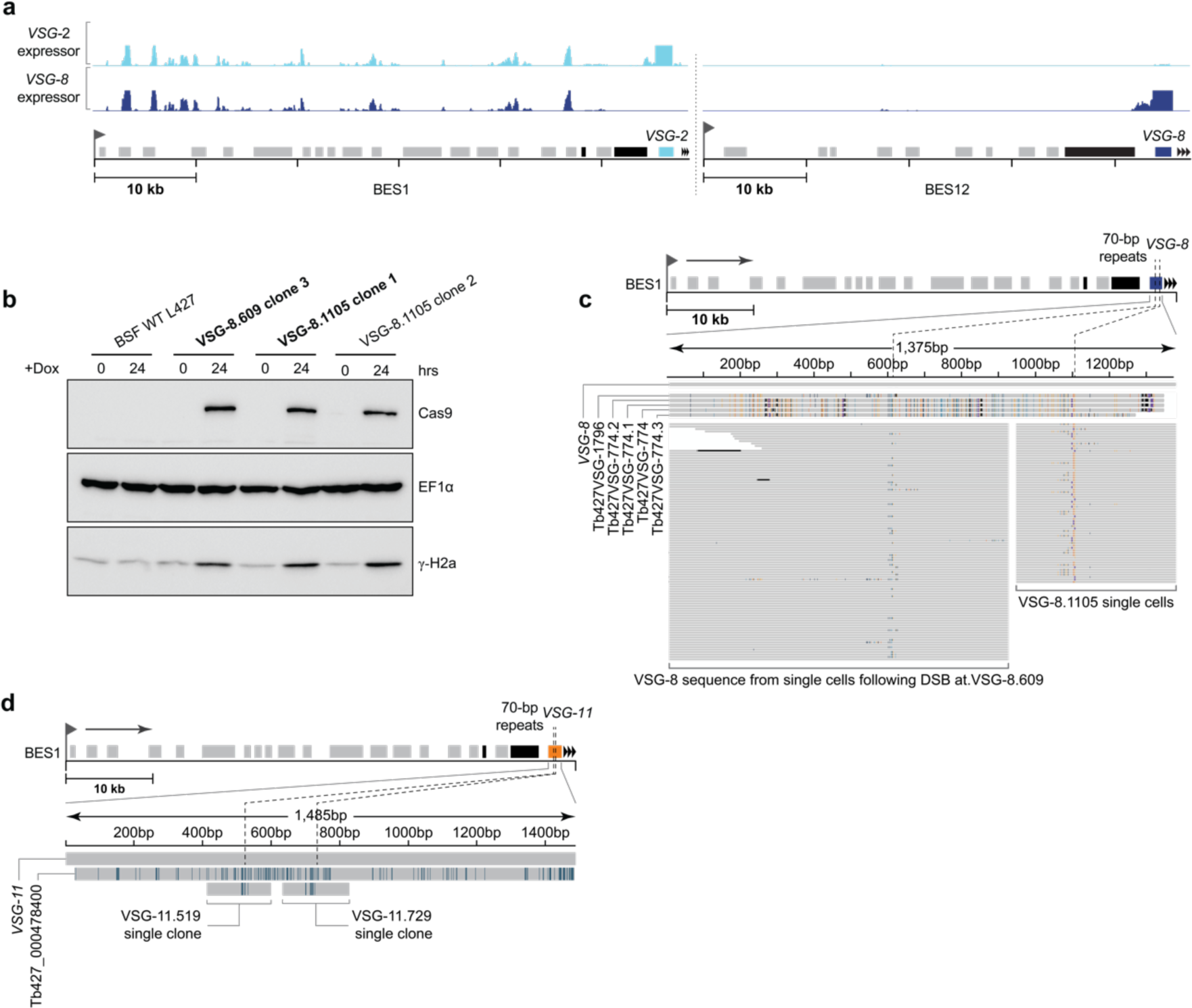
DSB in VSGs with homologous sequences in the genome lead to segmental gene conversion. **a,** Read coverage from bulk RNA-seq data on BES1 (left panel) and BES12 (right panel) for a control *VSG-2* expressing cell line and the *VSG-8* expressing cell line cloned from a DSB induction experiment on nucleotide position 152 of *VSG-2* CDS; showing that in this cell line VSG-8 was recombined and is being expressed from BES1. **b,** Western blotting showing Cas9 and γ-H2a expression levels before and 24 hours post-Cas9 induction with doxycycline (Dox) for two cell lines - VSG-8.609 (clone 3) and VSG-8.1105 (clones 1 and 2) - expressing different sgRNA for generating a DSB at positions 609 and 1105, respectively. Clones used for scRNA-seq experiments are in bold. **c,** *De novo* assembled VSG transcripts from individual cells after 96 hours DSB induction at nucleotide position 609 and 1105 of *VSG-8* CDS, aligned against *VSG-8* sequence. Five homologous VSGs genes or pseudogenes (potential ‘donors’) found in the *T. brucei* genome by BLAST search are also included on top of the alignment. Mismatched bases are colored based on the nucleotide identity. **d,** Sanger sequencing results for clones derived from a *VSG-11* expressing cell line after DSB induction at nucleotide positions 519 and 729 of *VSG-11* CDS. A single clone is shown for each cut. Sanger sequences are aligned against the reference *VSG-11* and a potential sequence ‘donor’ (Tb427_000478400:pseudogene, Tb427v11 genome).

**Extended Data Figure 6.**
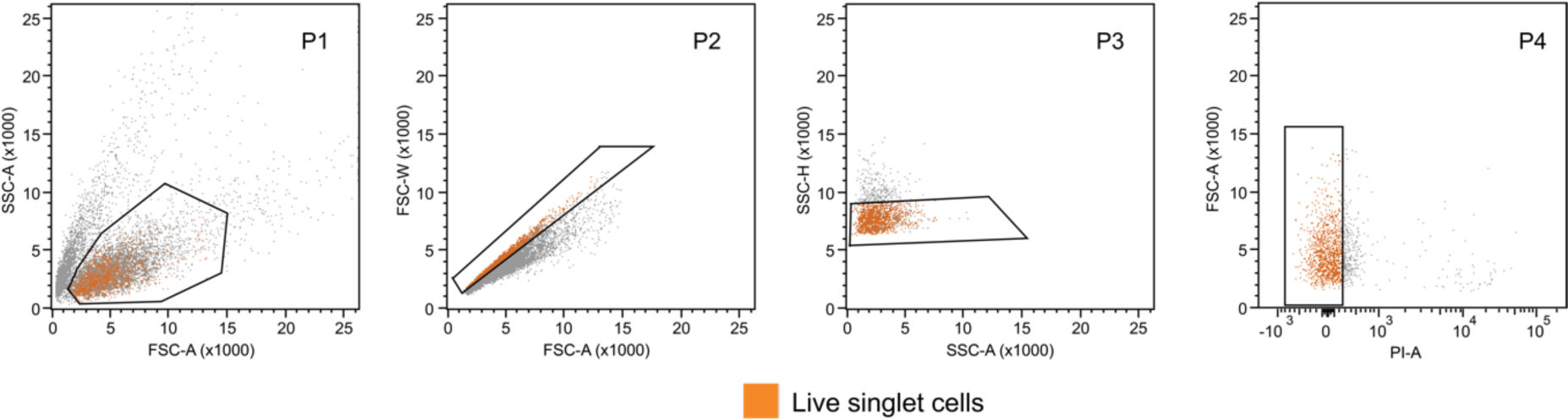
FACS gating strategy. FACS gating strategy for SL-Smart-seq3xpress library preparation. The cells were gated to exclude debris (P1), doublets (P2 and P3), and dead cells (P4). Propidium iodide was used to gate for live cells. The panel shows that P4 (colored orange) was used as a final cell population for sorting into the 384-well plates.

## References

1 Deitsch, K. W., Lukehart, S. A. & Stringer, J. R. Common strategies for antigenic variation by bacterial, fungal and protozoan pathogens. Nat Rev Microbiol 7, 493–503 (2009). 10.1038/nrmicro2145

2 Barcons-Simon, A., Carrington, M. & Siegel, T. N. Decoding the impact of nuclear organization on antigenic variation in parasites. Nat Microbiol 8, 1408–1418 (2023). 10.1038/s41564-023-01424-9

3 Trindade, S. et al. Trypanosoma brucei Parasites Occupy and Functionally Adapt to the Adipose Tissue in Mice. Cell Host Microbe 19, 837–848 (2016). 10.1016/j.chom.2016.05.002

4 Capewell, P. et al. The skin is a significant but overlooked anatomical reservoir for vector-borne African trypanosomes. Elife 5 (2016). 10.7554/eLife.17716

5 Cosentino, R. O., Brink, B. G. & Siegel, T. N. Allele-specific assembly of a eukaryotic genome corrects apparent frameshifts and reveals a lack of nonsense-mediated mRNA decay. NAR Genom Bioinform 3, lqab082 (2021). 10.1093/nargab/lqab082

6 Horn, D. & Barry, J. D. The central roles of telomeres and subtelomeres in antigenic variation in African trypanosomes. Chromosome Res 13, 525–533 (2005). 10.1007/s10577-005-0991-8

7 Wickstead, B., Ersfeld, K. & Gull, K. The small chromosomes of Trypanosoma brucei involved in antigenic variation are constructed around repetitive palindromes. Genome Res 14, 1014–1024 (2004). 10.1101/gr.2227704

8 Berriman, M. et al. The architecture of variant surface glycoprotein gene expression sites in Trypanosoma brucei. Mol Biochem Parasitol 122, 131–140 (2002).

9 Hertz-Fowler, C. et al. Telomeric expression sites are highly conserved in Trypanosoma brucei. PLoS One 3, e3527 (2008). 10.1371/journal.pone.0003527

10 Müller, L. S. M. et al. Genome organization and DNA accessibility control antigenic variation in trypanosomes. Nature 563, 121–125 (2018). 10.1038/s41586-018-0619-8

11 Van der Ploeg, L. H. & Cornelissen, A. W. The contribution of chromosomal translocations to antigenic variation in Trypanosoma brucei. Philos Trans R Soc Lond B Biol Sci 307, 13–26 (1984). 10.1098/rstb.1984.0105

12 Hoeijmakers, J. H., Frasch, A. C., Bernards, A., Borst, P. & Cross, G. A. Novel expression-linked copies of the genes for variant surface antigens in trypanosomes. Nature 284, 78–80 (1980). 10.1038/284078a0

13 Morrison, L. J., Majiwa, P., Read, A. F. & Barry, J. D. Probabilistic order in antigenic variation of Trypanosoma brucei. Int J Parasitol 35, 961–972 (2005). 10.1016/j.ijpara.2005.05.004

14 Mugnier, M. R., Cross, G. A. & Papavasiliou, F. N. The in vivo dynamics of antigenic variation in Trypanosoma brucei. Science 347, 1470–1473 (2015). 10.1126/science.aaa4502

15 Timmers, H. T., de Lange, T., Kooter, J. M. & Borst, P. Coincident multiple activations of the same surface antigen gene in Trypanosoma brucei. J Mol Biol 194, 81–90 (1987). 10.1016/0022-2836(87)90717-0

16 Lythgoe, K. A., Morrison, L. J., Read, A. F. & Barry, J. D. Parasite-intrinsic factors can explain ordered progression of trypanosome antigenic variation. Proc Natl Acad Sci U S A 104, 8095–8100 (2007). 10.1073/pnas.0606206104

17 Gjini, E., Haydon, D. T., Barry, J. D. & Cobbold, C. A. Critical Interplay between Parasite Differentiation, Host Immunity, and Antigenic Variation in Trypanosome Infections. The American Naturalist 176, 424–439 (2010). 10.1086/656276

18 Liu, D., Albergante, L., Newman, T. J. & Horn, D. Faster growth with shorter antigens can explain a VSG hierarchy during African trypanosome infections: a feint attack by parasites. Sci Rep 8, 10922 (2018). 10.1038/s41598-018-29296-8

19 Escrivani, D. O., Scheidt, V., Tinti, M., Faria, J. & Horn, D. Competition among variants is predictable and contributes to the antigenic variation dynamics of African trypanosomes. PLoS Pathog 19, e1011530 (2023). 10.1371/journal.ppat.1011530

20 Hagemann-Jensen, M., Ziegenhain, C. & Sandberg, R. Scalable single-cell RNA sequencing from full transcripts with Smart-seq3xpress. Nat Biotechnol 40, 1452–1457 (2022). 10.1038/s41587-022-01311-4

21 Glover, L., Alsford, S. & Horn, D. DNA break site at fragile subtelomeres determines probability and mechanism of antigenic variation in African trypanosomes. PLoS Pathog 9, e1003260 (2013). 10.1371/journal.ppat.1003260

22 Robinson, N. P., Burman, N., Melville, S. E. & Barry, J. D. Predominance of duplicative VSG gene conversion in antigenic variation in African trypanosomes. Mol Cell Biol 19, 5839–5846 (1999).

23 Islam, S. et al. Quantitative single-cell RNA-seq with unique molecular identifiers. Nat Methods 11, 163–166 (2014). 10.1038/nmeth.2772

24 Haanstra, J. R. et al. Control and regulation of gene expression: quantitative analysis of the expression of phosphoglycerate kinase in bloodstream form Trypanosoma brucei. J Biol Chem 283, 2495–2507 (2008). 10.1074/jbc.M705782200

25 Hagemann-Jensen, M. et al. Single-cell RNA counting at allele and isoform resolution using Smart-seq3. Nat Biotechnol 38, 708–714 (2020). 10.1038/s41587-020-0497-0

26 Sutton, R. E. & Boothroyd, J. C. Evidence for trans splicing in trypanosomes. Cell 47, 527–535 (1986). 10.1016/0092-8674(86)90617-3

27 Sinha, R. et al. Index switching causes “spreading-of-signal” among multiplexed samples in Illumina HiSeq 4000 DNA sequencing. bioRxiv, 125724 (2017). 10.1101/125724

28 Briggs, E. M., Rojas, F., McCulloch, R., Matthews, K. R. & Otto, T. D. Single-cell transcriptomic analysis of bloodstream Trypanosoma brucei reconstructs cell cycle progression and developmental quorum sensing. Nat Commun 12, 5268 (2021). 10.1038/s41467-021-25607-2

29 Ding, J. et al. Systematic comparison of single-cell and single-nucleus RNA-sequencing methods. Nat Biotechnol 38, 737–746 (2020). 10.1038/s41587-020-0465-8

30 Faria, J. et al. Spatial integration of transcription and splicing in a dedicated compartment sustains monogenic antigen expression in African trypanosomes. Nat Microbiol 6, 289–300 (2021). 10.1038/s41564-020-00833-4

31 Figueiredo, L. M., Janzen, C. J. & Cross, G. A. A histone methyltransferase modulates antigenic variation in African trypanosomes. PLoS Biol 6, e161 (2008). 10.1371/journal.pbio.0060161

32 Faria, J. R. C. et al. An allele-selective inter-chromosomal protein bridge supports monogenic antigen expression in the African trypanosome. Nat Commun 14, 8200 (2023). 10.1038/s41467-023-44043-y

33 Muskovic, W. & Powell, J. E. DropletQC: improved identification of empty droplets and damaged cells in single-cell RNA-seq data. Genome Biol 22, 329 (2021). 10.1186/s13059-021-02547-0

34 Thivolle, A. et al. DNA double strand break position leads to distinct gene expression changes and regulates VSG switching pathway choice. PLoS Pathog 17, e1010038 (2021). 10.1371/journal.ppat.1010038

35 Glover, L., McCulloch, R. & Horn, D. Sequence homology and microhomology dominate chromosomal double-strand break repair in African trypanosomes. Nucleic Acids Res 36, 2608–2618 (2008). 10.1093/nar/gkn104

36 Hovel-Miner, G., Mugnier, M. R., Goldwater, B., Cross, G. A. & Papavasiliou, F. N. A Conserved DNA Repeat Promotes Selection of a Diverse Repertoire of Trypanosoma brucei Surface Antigens from the Genomic Archive. PLoS Genet 12, e1005994 (2016). 10.1371/journal.pgen.1005994

37 Burton, P., McBride, D. J., Wilkes, J. M., Barry, J. D. & McCulloch, R. Ku heterodimer-independent end joining in Trypanosoma brucei cell extracts relies upon sequence microhomology. Eukaryot Cell 6, 1773–1781 (2007). 10.1128/ec.00212-07

38 Aresta-Branco, F., Pimenta, S. & Figueiredo, L. M. A transcription-independent epigenetic mechanism is associated with antigenic switching in Trypanosoma brucei. Nucleic Acids Res 44, 3131–3146 (2016). 10.1093/nar/gkv1459

39 Hall, J. P., Wang, H. & Barry, J. D. Mosaic VSGs and the scale of Trypanosoma brucei antigenic variation. PLoS Pathog 9, e1003502 (2013). 10.1371/journal.ppat.1003502

40 Briggs, E., Crouch, K., Lemgruber, L., Lapsley, C. & McCulloch, R. Ribonuclease H1-targeted R-loops in surface antigen gene expression sites can direct trypanosome immune evasion. PLoS Genet 14, e1007729 (2018). 10.1371/journal.pgen.1007729

41 Girasol, M. J. et al. RAD51-mediated R-loop formation acts to repair transcription-associated DNA breaks driving antigenic variation in Trypanosoma brucei. Proc Natl Acad Sci U S A 120, e2309306120 (2023). 10.1073/pnas.2309306120

42 Alsford, S., Kawahara, T., Glover, L. & Horn, D. Tagging a T. brucei RRNA locus improves stable transfection efficiency and circumvents inducible expression position effects. Mol Biochem Parasitol 144, 142–148 (2005). 10.1016/j.molbiopara.2005.08.009

43 Rico, E., Jeacock, L., Kovarova, J. & Horn, D. Inducible high-efficiency CRISPR-Cas9-targeted gene editing and precision base editing in African trypanosomes. Sci Rep 8, 7960 (2018). 10.1038/s41598-018-26303-w

44 Hirumi, H. & Hirumi, K. Continuous cultivation of Trypanosoma brucei blood stream forms in a medium containing a low concentration of serum protein without feeder cell layers. J Parasitol 75, 985–989 (1989).

45 Burkard, G., Fragoso, C. M. & Roditi, I. Highly efficient stable transformation of bloodstream forms of Trypanosoma brucei. Mol Biochem Parasitol 153, 220–223 (2007). 10.1016/j.molbiopara.2007.02.008

46 MacPherson, C. R. & Scherf, A. Flexible guide-RNA design for CRISPR applications using Protospacer Workbench. Nat Biotechnol 33, 805–806 (2015). 10.1038/nbt.3291

47 Dejung, M. et al. Quantitative Proteomics Uncovers Novel Factors Involved in Developmental Differentiation of Trypanosoma brucei. PLoS Pathog 12, e1005439 (2016). 10.1371/journal.ppat.1005439

48 Pinger, J., Chowdhury, S. & Papavasiliou, F. N. Variant surface glycoprotein density defines an immune evasion threshold for African trypanosomes undergoing antigenic variation. Nat Commun 8, 828 (2017). 10.1038/s41467-017-00959-w

49 Martin, M. Cutadapt removes adapter sequences from high-throughput sequencing reads. 2011 17, 3 (2011). 10.14806/ej.17.1.200

50 Li, H. seqtk: Toolkit for processing sequences in FASTA/Q formats, <https://github.com/lh3/seqtk>

51 Dobin, A. et al. STAR: ultrafast universal RNA-seq aligner. Bioinformatics 29, 15–21 (2013). 10.1093/bioinformatics/bts635

52 Kaminow, B., Yunusov, D. & Dobin, A. STARsolo: accurate, fast and versatile mapping/quantification of single-cell and single-nucleus RNA-seq data. bioRxiv, 2021.2005.2005.442755 (2021). 10.1101/2021.05.05.442755

53 Larsson, A. J. M., Stanley, G., Sinha, R., Weissman, I. L. & Sandberg, R. Computational correction of index switching in multiplexed sequencing libraries. Nat Methods 15, 305–307 (2018). 10.1038/nmeth.4666

54 Ramírez, F., Dündar, F., Diehl, S., Grüning, B. A. & Manke, T. deepTools: a flexible platform for exploring deep-sequencing data. Nucleic Acids Res 42, W187–191 (2014). 10.1093/nar/gku365

55 Lopez-Delisle, L. et al. pyGenomeTracks: reproducible plots for multivariate genomic datasets. Bioinformatics 37, 422–423 (2021). 10.1093/bioinformatics/btaa692

56 Robinson, J. T. et al. in Nat Biotechnol Vol. 29 24–26 (2011).

57 Shanmugasundram, A. et al. TriTrypDB: An integrated functional genomics resource for kinetoplastida. PLoS Negl Trop Dis 17, e0011058 (2023). 10.1371/journal.pntd.0011058

58 Renaud, G., Stenzel, U., Maricic, T., Wiebe, V. & Kelso, J. deML: robust demultiplexing of Illumina sequences using a likelihood-based approach. Bioinformatics 31, 770–772 (2015). 10.1093/bioinformatics/btu719

59 Grabherr, M. G. et al. Full-length transcriptome assembly from RNA-Seq data without a reference genome. Nat Biotechnol 29, 644–652 (2011). 10.1038/nbt.1883

60 Camacho, C. et al. BLAST+: architecture and applications. BMC Bioinformatics 10, 421 (2009). 10.1186/1471-2105-10-421

61 Li, H. et al. The Sequence Alignment/Map format and SAMtools. Bioinformatics 25, 2078–2079 (2009). 10.1093/bioinformatics/btp352

